# Fluctuations in TCR and pMHC interactions regulate T cell activation

**DOI:** 10.1101/2021.02.09.430441

**Authors:** Joseph R. Egan, Enas Abu-Shah, Omer Dushek, Tim Elliott, Ben D. MacArthur

**Affiliations:** Mathematical Sciences, University of Southampton, Southampton SO17 1BJ, United Kingdom; Centre for Cancer Immunology, University Hospital Southampton, Southampton SO16 6YD, United Kingdom; Institute for Life Sciences, University of Southampton, Southampton SO17 1BJ, United Kingdom; Sir William Dunn School of Pathology, University of Oxford, Oxford OX1 3RE, United Kingdom; Kennedy Institute of Rheumatology, University of Oxford, Oxford OX3 7FY, United Kingdom; Nuffield Department of Medicine, University of Oxford, Oxford OX3 7BN, United Kingdom; Centre for Human Development, Stem Cells and Regeneration, University of Southampton, Southampton SO17 1BJ, United Kingdom; Alan Turing Institute, London NW1 2DB, United Kingdom

## Abstract

Adaptive immune responses depend on interactions between T cell receptors (TCRs) and peptide major-histocompatibility complex (pMHC) ligands located on the surface of T cells and antigen presenting cells (APCs) respectively. As TCRs and pMHCs are often only present at low copy-numbers their interactions are inherently stochastic, yet the role of stochastic fluctuations on T cell function is unclear. Here we introduce a minimal stochastic model of T cell activation that accounts for serial TCR-pMHC engagement, reversible TCR conformational change and TCR aggregation. Analysis of this model indicates that it is not the strength of binding between the T cell and the APC cell *per se* that elicits an immune response, but rather the information imparted to the T cell from the encounter, as assessed by the entropy rate of the TCR-pMHC binding dynamics. This view provides an information-theoretic interpretation of T cell activation that explains a range of experimental observations. Based on this analysis we propose that effective T cell therapeutics may be enhanced by optimizing the inherent stochasticity of TCR-pMHC binding dynamics.

## Introduction

Lymphocytes are responsible for immunity and a subset known as T cells are critical for adaptive immunity [1]. T cell receptors (TCRs) located on the T cell surface reversibly bind to peptide-major histocompatibility complex (pMHC) ligands located on the surface of antigen presenting cells (APCs) [2]. This interaction can generate a signaling cascade within the T cell [3] leading to a variety of functional responses [4], including the production of soluble messengers called cytokines [5]. Furthermore, an activated T cell is stimulated to proliferate, thereby generating progeny that can differentiate into effector cells [3]. These mature T cells are then able to clear antigen from the body by seeking out and destroying harmful pathogen-infected or tumor cells [6]. Yet despite decades of research, it is still unclear which TCR proximal mechanisms are primarily responsible for transmitting the information encoded in the pMHC ligand to the T cell intracellular signaling pathways [7, 8, 9, 10, 11, 12, 13, 14].

Each TCR has a short intracellular domain which, alone, does not have the capacity to initiate signaling [1]. Consequently, a TCR associates with three CD3 subunits to facilitate signal transduction to the T cell interior [15]. The CD3 subunits have tails extending into the cytoplasm that contain multiple copies of the immuno-receptor tyrosine activation motif (ITAM) [3].

The phosphorylation of ITAMs is considered one of the earliest events in the signaling cascade that leads to T cell activation [4]. Kinases, such as LCK, are molecules that phosphorylate ITAMs and therefore favour signaling. In contrast, phosphatases, such as CD45, are molecules that dephosphorylate ITAMs and therefore inhibit signaling.

Three to four main mechanisms have been proposed to initiate signaling events following pMHC-TCR binding [16, 15, 2, 6, 17, 18, 14], all of which are likely to shift the balance in favour of ITAM phosphorylation [15]. One mechanism is the segregation of CD45 molecules from the TCR-CD3 complex [16] that could allow for the stable phosphorylation of ITAMs by LCK. A second mechanism is the aggregation of TCR-CD3 complexes and their subsequent ‘microcluster’ formation [19, 20, 21] that could increase the proximity of LCK molecules leading to enhanced ITAM phosphorylation. A third mechanism is a physical and/or chemical change (generally referred to as a conformational change [12]) in the TCR-CD3 complex, possibly in the cytoplasmic tails that could expose their ITAMs to enhanced phosphorylation. A fourth mechanism [14] (which is arguably a sub-mechanism of the third mechanism [15]) is the generation of forces tangential to the T cell surface caused by the movement of T cells as they scan the surface of APCs for antigenic peptides [13]. These mechanical forces, such as pulling or shearing, could lead to the uncoupling of the CD3 tails from the T cell membrane, exposing their ITAMs to phosphorylation by LCK.

It is likely that not one mechanism alone is responsible for the initiation of signaling events [22]. For example, it has been proposed that mechanical forces induce conformational changes [15, 2, 23, 13, 14] which subsequently induce aggregation and clustering [15, 17]. Others have advocated that conformational change is instead directly induced by pMHC ligand binding [24, 25] and also reversible [26, 7, 27, 18, 14]. It has also been argued that both conformational TCR change and TCR clustering are necessary for T cell activation [8] and may improve antigen discrimination [28, 29]. Although some have argued that it is unnecessary [30, 18, 31], others have supported the view that signaling requires just a few pMHC ligands to serially bind multiple TCRs [32, 33, 34, 35] and that this serial engagement could lead to a conformational change in each TCR [36, 37]. The three mechanisms of serial TCR-pMHC engagement; reversible TCR conformational change and TCR aggregation are shown schematically in Fig. 1. Notably, it has been suggested that a combination of these three mechanisms may allow the T cell to efficiently scan the APC surface with high specificity and sensitivity for rare pMHC ligands presented at low copy-numbers [32, 38, 15, 1, 10, 4, 35].

**Figure 1.**
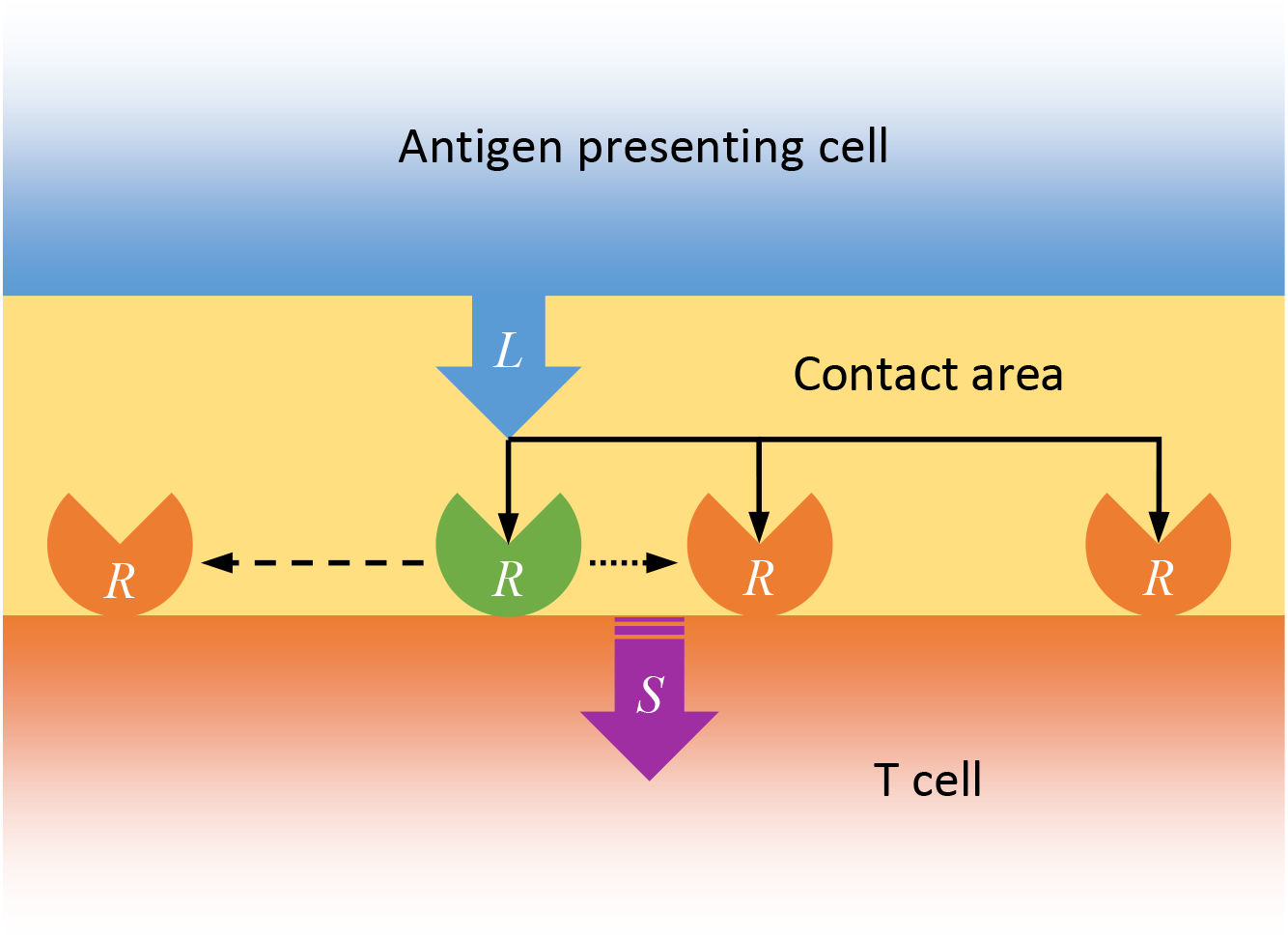
Schematic of the three modeled mechanisms involved in TCR-pMHC binding. (1) Solid black arrows represent a pMHC ligand, *L* serially engaging with multiple TCRs, *R* within the contact area. (2) The green TCR represents a conformational change upon pMHC ligand binding. The dashed black arrow represents the TCR reverting back to its original state at some time after unbinding. (3) The dotted black arrow represents TCR aggregation following pMHC ligand binding. The combination of these thee mechanisms generates a signal, *S* within the T cell.

Indeed, there is increasing evidence that T cell activation can be induced by as few as ~1-10 pMHC ligands [39, 40, 41, 33, 30, 35] and that microclusters may contain as few as ~10-100 TCRs [19, 20, 21, 42, 32, 33]. At such low copy-numbers the TCR-pMHC binding dynamics are inherently stochastic, yet the effect of this stochasticity on T cell activation is unclear. Stochastic fluctuations have been shown to be functionally important in numerous other biological contexts [43, 44, 45] and, therefore, it is conceivable that the T cell has evolved to utilize these fluctuations to enhance its own function.

Here, we develop a minimal stochastic model of the TCR-pMHC binding dynamics that includes serial TCR-pMHC engagement, reversible TCR conformational change and TCR aggregation. We show that, collectively, these three mechanisms are both necessary and sufficient for the T cell to convert stochastic fluctuations in the TCR-pMHC binding dynamics into a well-defined signal. Based on this analysis we propose that the T cell response to an APC is not determined by the strength of TCR-pMHC binding *per se*, but rather by the information conveyed to the T cell by the encounter, as assessed by the entropy rate of the TCR-pMHC binding dynamics. We validate this hypothesis against a range of experimental studies, including a number of dose-response data-sets, before discussing the implications for T cell based therapeutics.

## Results

### Fluctuations in TCR-pMHC binding dynamics generate information

To start, we will introduce some information-theoretic notions in the context of a simple model of TCR-pMHC binding, before discussing how they apply to a more realistic model of T cell activation.

Consider the process of TCR-pMHC reversible heterodimerization, given by the following reactions:

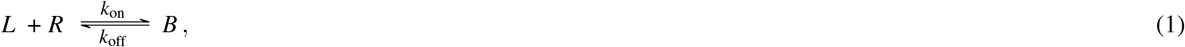

where *L* denotes the pMHC ligand, *R* denotes the TCR, *B* denotes the TCR-pMHC complex, *k*_off_ is the rate of unbinding, *k*_on_/*ν* is the rate of binding [46] and *ν* is the 2-dimensional (2D) contact area in which the biochemical reactions take place.

At low copy-numbers these reactions will be inherently stochastic and the copy-number of the TCR-pMHC complex will accordingly fluctuate randomly over time. To quantify the extent of this stochasticity we will use two measures. First, the Shannon entropy, *H*(*B*) (in bits) given by:

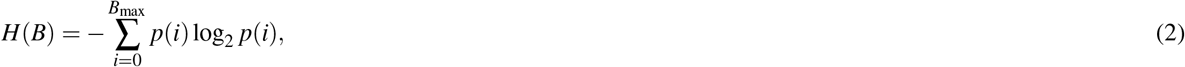

where *B*_max_ is the maximum number of TCR-pMHC complexes (given by Eq. 11 in the **Materials and Methods**) and *p*(*i*) is the stationary probability that *i* copies of the TCR-pMHC complex are present (given by Eq. 13 in the **Materials and Methods**). In what follows, we will assume that the T cell responds on a slower timescale than the TCR-pMHC binding dynamics, and consider properties of stationary probability distributions only. In general, the Shannon entropy is a simple measure of information or ‘disorder’ [47]. In the context of the T cell-APC contact area, it is the average amount of information imparted to the T cell per TCR-pMHC binding/unbinding event. As such, although it is a useful measure of information, the Shannon entropy does not take account of the speed of the underlying reactions which will vary with the kinetic rate parameters. Therefore, the Shannon entropy cannot distinguish between fast and slow dynamics.

To clarify this distinction we will use an alternative measure: the entropy rate, *H*′(*B*) (in bits per second) which is calculated as the mean reaction rate (i.e. the average number of binding/unbinding events per second) multiplied by the Shannon entropy. For the reversible heterodimerization reactions given in Eq. 1, the entropy rate is:

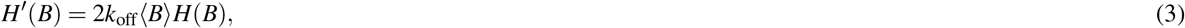

where (*B*) is the mean of the TCR-pMHC complex stationary probability distribution (for details see section 2.2 of the **Supplementary Information**). In the context of the T cell-APC contact area, the entropy rate is the average amount of information imparted to the T cell by the TCR-pMHC binding dynamics per second. Therefore, unlike the Shannon entropy, the entropy rate can distinguish between fast and slow dynamics.

While it has a useful information-theoretic interpretation, the entropy rate is complex to calculate in practice. However, we can similarly define the ‘variance rate’, Var′(*B*) as:

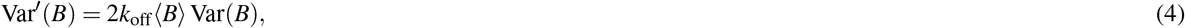

where Var(*B*) is the variance of the TCR-pMHC complex stationary probability distribution. Although the variance rate does not have an information theoretic interpretation, it exhibits similar features to the entropy rate for the simple dynamics described here and is more analytically tractable (for details see sections 2.3 and 4.2 of the Supplementary Information). We will make use of this connection in the next section, where we analyze a more realistic model of TCR-pMHC binding dynamics.

To illustrate these concepts, **Fig. 2** shows some representative stochastic simulations of TCR-pMHC reversible heterodimerization using Gillespie’s direct method [48, 49]. Three features of these simulations are notable.

**Figure 2.**
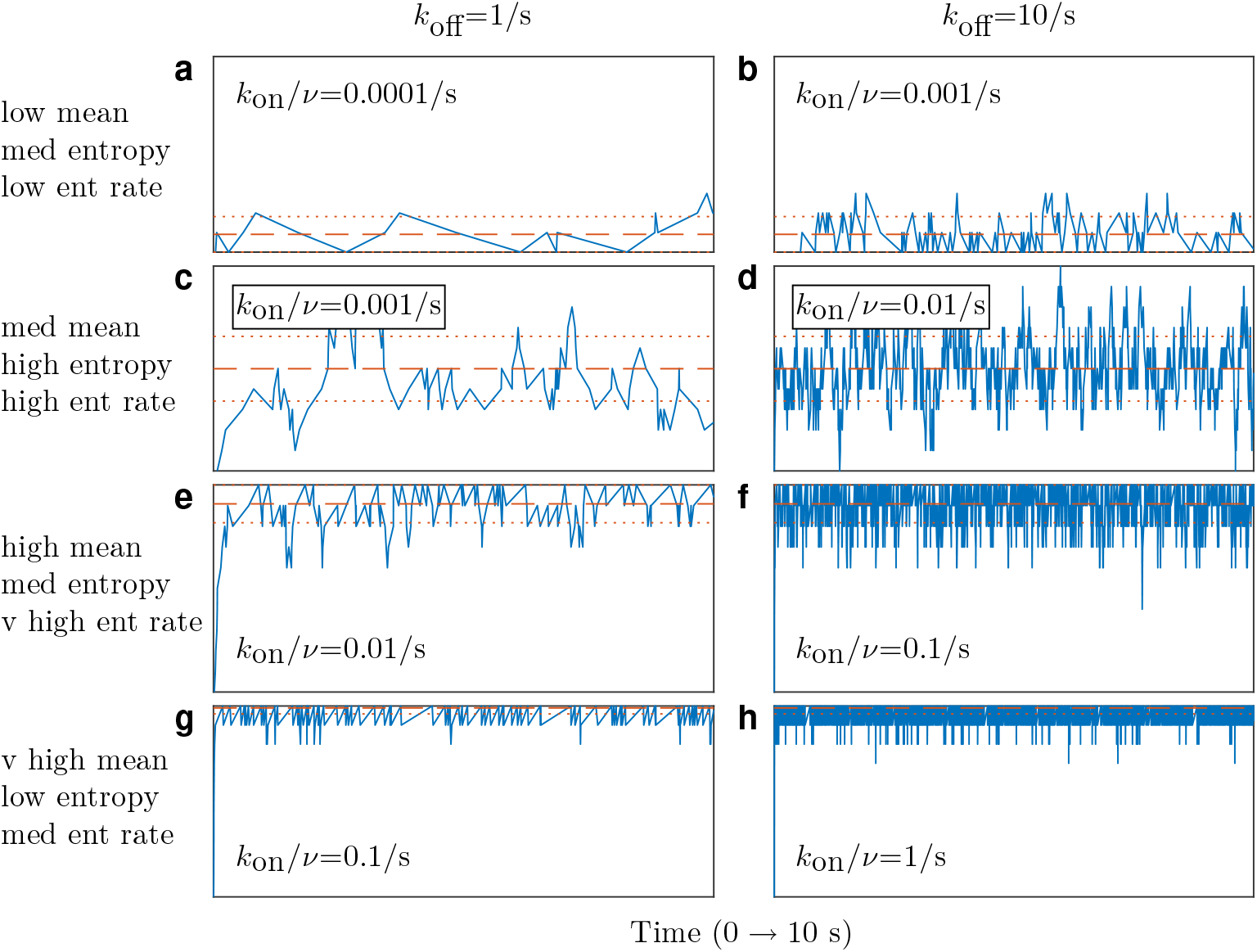
Fluctuations in TCR-pMHC dynamics generate information. Blue lines show representative stochastic simulations of the TCR-pMHC complex copy-number, *B* for the first 10 seconds of the reversible heterodimerization reactions given in Eq. 1. Dashed red lines show the mean and dotted red lines show the mean plus/minus one standard deviation. In all panels the number of TCRs, *R*_max_ = 10 and the number of pMHC ligands, *L*_max_ = 1, 000 which gives the maximum number of TCR-pMHC complexes, *B*_max_ = 10 via Eq. 11. The binding rate, *k*_on_/*ν* and unbinding rate, *k*_off_ are varied over orders of magnitude within a plausible physiological range as described in the **Materials and Methods**.

First, while the mean number of TCR-pMHC complexes increases monotonically with the binding rate (*k*_on_/*ν*), both the Shannon entropy and the entropy rate initially increase as the binding rate increases, but then decrease as the binding rate increases further still (to see this compare the panels in each column of **Fig. 2**). This biphasic pattern occurs because fluctuations are minimal when binding is very weak or very strong (i.e., when complexes do not easily associate or dissociate respectively) yet become larger at intermediate affinities that allow both binding and unbinding events to easily occur. Second, while the mean number of TCR-pMHC complexes and Shannon entropy are dependent on three model parameters: the total number of pMHC ligands and TCRs at the contact area (which we denote *L*_max_ and *R*_max_ respectively) and the 2D dissociation constant, *K*_d_ given by:

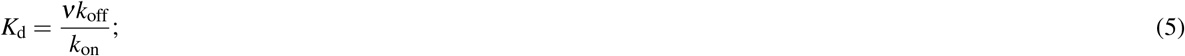

the entropy rate is explicitly dependent on both the binding rate and the unbinding rate (rather than simply the ratio of the two, *K*_d_). Thus, dynamics associated with different kinetic rate parameters may have the same mean number of TCR-pMHC complexes and Shannon entropy, but very different entropy rates (to see this compare the panels in each row of **Fig. 2**. In each case, the entropy rate in the right column is an order of magnitude higher than in the left column).

Third, for a fixed unbinding rate, the maximum entropy rate (and therefore the maximum rate at which information can be imparted to the T cell) is achieved via a trade-off between the average number of TCR-pMHC complexes and average magnitude of the stochastic fluctuations. So, the TCR-pMHC binding dynamics illustrated in **Fig. 2f** have the largest entropy rate of all the panels because they combine both a relatively high mean with a relatively high Shannon entropy.

Collectively, this reasoning suggests that fluctuations in TCR-pMHC binding dynamics can generate information and thereby may have an important, but as yet unexplored, part to play in regulating T cell activation.

### TCR-pMHC fluctuations regulate T cell activation

To investigate this possibility further we sought to construct a minimal model of the TCR-pMHC binding dynamics that includes the effects of serial TCR engagement, reversible TCR conformational change and TCR aggregation. Our minimal model consists of the following set of reactions:

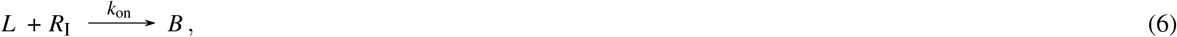

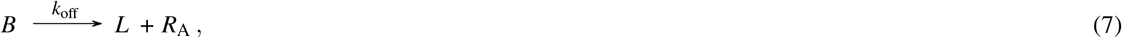

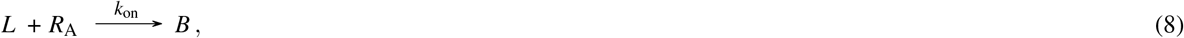

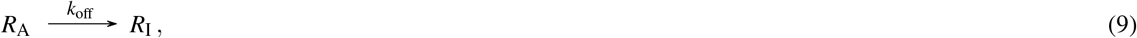

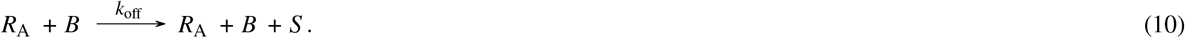

where *R*_I_ and *R*_A_, denote ‘inactive’ TCRs and ‘active’ TCRs respectively, and *S* denotes an activating T cell signal. Note that an active TCR can be interpreted as one that has undergone a conformational change due to pMHC ligand binding and the generation of a signal can be interpreted as a subsequent consequence of TCR aggregation; see **Fig. 1**. We emphasize that Eqs. 6–10 are not meant to be a detailed model of every aspect of TCR-pMHC binding and T cell activation. Rather, they encapsulate key mechanisms in a parsimonious way that allows for a transparent exploration of their consequences. Particularly, Eq. 10 captures salient features of TCR aggregation without recourse to relatively complex stochastic reaction-diffusion processes which, although more mechanistically detailed, may be less tractable and harder to interpret. A more detailed explanation of how each reaction relates to each of the three TCR proximal mechanisms detailed in **Fig. 1** is provided in the **Materials and Methods**.

This modeling framework is useful because it accounts for additional mechanisms of importance, yet central aspects of the reversible heterodimerization reactions given in Eq. 1 are conserved (for details see section 5 of the **Supplementary Information**). In particular, the dynamics of the TCR-pMHC complex number, *B*, pMHC ligand number, *L*, and the sum of the inactive and active TCR numbers, *R* = *R*_I_ + *R*_A_, are equivalent to those of the straightforward reversible heterodimerization reactions. Thus, calculations of the mean number of TCR-pMHC complexes, Shannon entropy, and the variance/entropy rates described in the previous section also apply to this model.

Moreover, the effects of these quantities on signal generation may now be explored. In section 5.1 of the the **Supplementary Information** we show that the mean number of active TCRs, ⟨*R*_A_⟩, is equal to the variance of the TCR-pMHC complex number, Var(*B*), in wide regions of parameter space. This is notable because in this framework a T cell signal is stochastically generated if an active TCR is in close proximity to a TCR-pMHC complex (see Eq. 10). Consequently, this implies that the mean signaling rate, 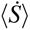 (i.e. the average rate at which a signal is generated) is approximately equal to half the variance rate, Var′(*B*) for a wide range of parameter values, as shown in **Fig. 3**. This reasoning suggests that serial TCR-pMHC engagement, reversible TCR conformational change and TCR aggregation work collectively to allow the T cell to process environmental information appropriately.

**Figure 3.**
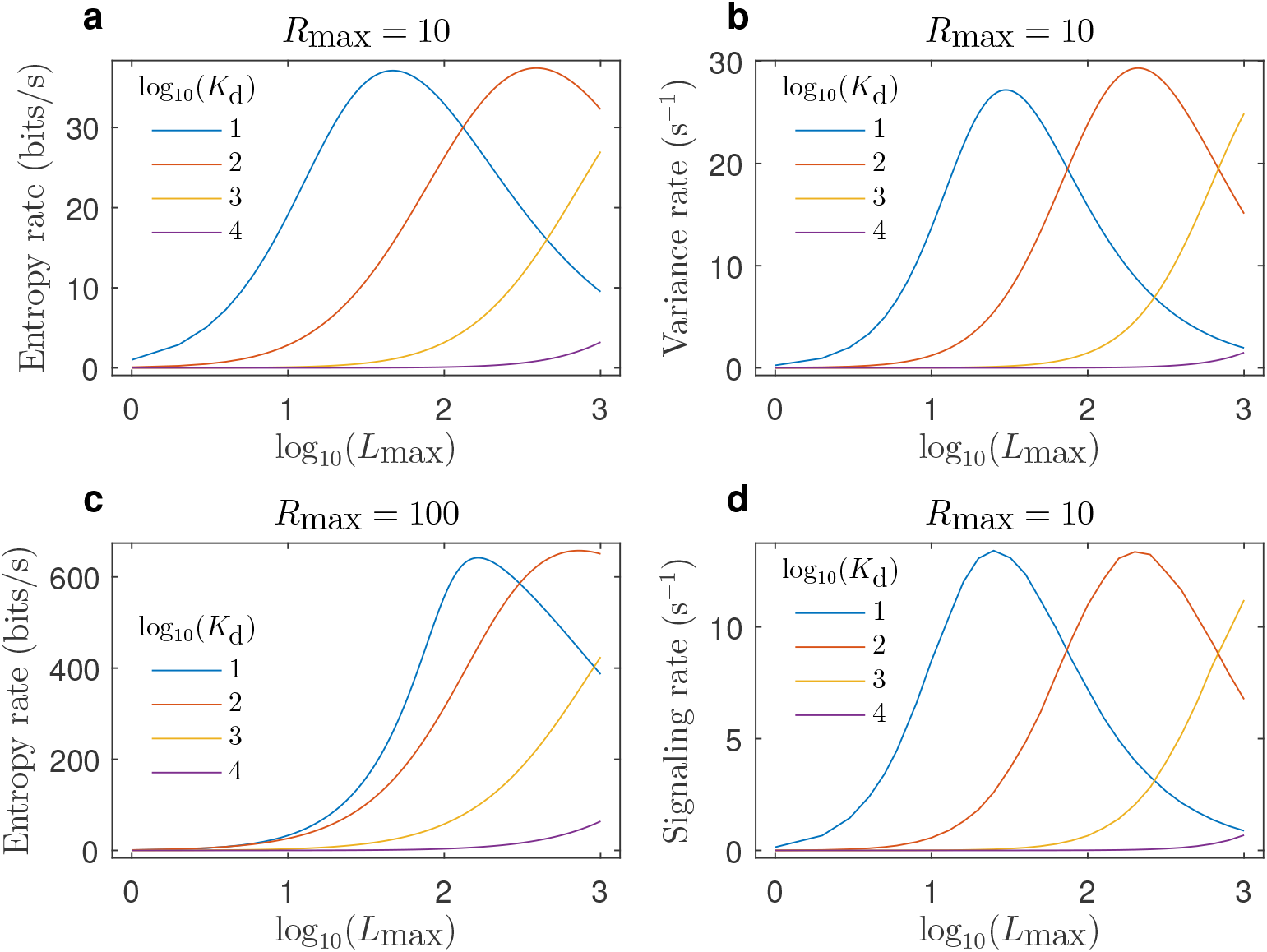
T cell signaling rate is regulated by TCR-pMHC fluctuations. Theoretical dose-response curves where dose on the x-axis is given by the number of pMHC ligands (*L*_max_) and the legend gives the 2D dissociation constant (*K*_d_). Response on the y-axis is given by the entropy rate (**a,c**) and variance rate (**b**) of the TCR-pMHC complex stationary probability distribution, and the mean signaling rate (**d**) based on the minimal model of T cell activation as calculated via stochastic simulations of the reactions given in Eqs. 6–10. Comparison of panels **a** and **c** shows that variation in the number of TCRs (*R*_max_) does not affect the qualitative nature of the curves. In all panels *k*_off_ = 1/*s*. See **Materials and Methods** for justification of these and other literature-derived parameter values.

In addition to offering an explanation of why these mechanisms are central to T cell activation, this perspective has important implications for optimization of the T cell response. In this minimal model, increasing the TCR-pMHC binding rate or the number of pMHC ligands (i.e. dose) initially increases the variance of the TCR-pMHC complex number and thereby the mean number of active TCRs and mean signaling rate (starting from a low binding rate or low dose). However, as the binding rate or dose increases further still, the variance of the TCR-pMHC complex number will fall and, while the mean number of TCR-pMHC complexes will continue to increase, the mean number of active TCRs will start to decrease. This, in turn, leads to a decrease in the mean signaling rate. Thus, because T cell activation is not regulated solely by the binding strength (i.e. affinity) between the TCR and pMHC molecules, but also by their dynamic fluctuations, maximal T cell activation is predicted to occur at an intermediate affinity (particularly with an intermediate to high physiological dose) or an intermediate dose (particularly with an intermediate to high physiological affinity). **Fig. 3** shows results of the binding and activation dynamics that illustrate these points, which are discussed further in the following section.

In section 5.2 of the **Supplementary Information** we show that a similar alternative minimal model gives a mean signaling rate that is equivalent to that shown in **Fig. 3d** over a wide area of parameter space. This suggests that the minimal model described by Eqs. 6–10 is representative of a wider class of models that exhibit shared characteristics. Two key characteristics that are common to both minimal models are that (1) TCRs are initially inactive and can become active following pMHC ligand binding and (2) the generation of a signal requires an unbound TCR and a bound TCR, one of which must be in an active state.

Collectively, these results indicate that stochastic fluctuations, as quantified by the variance rate of the TCR-pMHC binding dynamics, may regulate T cell activation. While we could not obtain a corresponding analytical result for the entropy rate, the variance rate and entropy rate are very closely related and numerical simulations of TCR-pMHC binding dynamics indicate a similar dependency (*cf* **Fig. 3a** and **Fig. 3b**). This suggests that the variance rate is an analytically convenient proxy for the more biologically-meaningful entropy rate as a measure of the magnitude and rate of TCR-pMHC fluctuations.

### Experimental validation

As described above, our model suggests that intermediate affinity and intermediate dose scenarios can give rise to highly stochastic, information-rich, dynamics which the T cell is able to process, via simple molecular mechanisms, into a defined cellular response. To investigate the validity of this view we sought to determine its experimental support.

First, a number of experimental studies have reported that the T cell response is maximized at an intermediate affinity [50, 51, 52, 53, 54, 55, 56, 57, 58, 59]. Moreover, some of these studies have shown that an optimal affinity exists for both proximal and distal activation events (i.e. early and later T cell responses) [54, 55, 56].

Second, other experimental studies have reported the T cell response is maximised at an intermediate dose [60, 61, 62, 63], particularly for higher affinity pMHC ligands [64, 65, 53, 66, 67, 68]. For example, **Figs. 4a-c** show dose-response curves from three previous studies [64, 65, 53] and **Fig. 4d** shows previously unpublished experimental dose-response data. Briefly, two of these studies, as well as our study, utilized a TCR and varied the affinity via a panel of ligands [64, 53]. The other study utilized a chimeric antigen receptor (CAR) and varied the affinity via its single-chain fragment binding domain [65]. In the two earlier studies the ligands were antibodies [64] or ErbB2 proteins [65] that were immobilized on micro-titer plates before being stimulated by T cells. In the later study and here, the ligands were altered peptide ligands that were either directly added to cultures containing T cells [53] or pulsed into monocyte-derived dendritic cells before being co-cultured with T cells (see **Materials and Methods**). Despite the experimental variation, all of the panels of **Fig. 4** are qualitatively consistent with our model predictions shown in **Fig. 3**. Furthermore, all of these data-sets exhibit similar features regardless of the T cell responses that were measured or the experimental techniques that were implemented.

**Figure 4.**
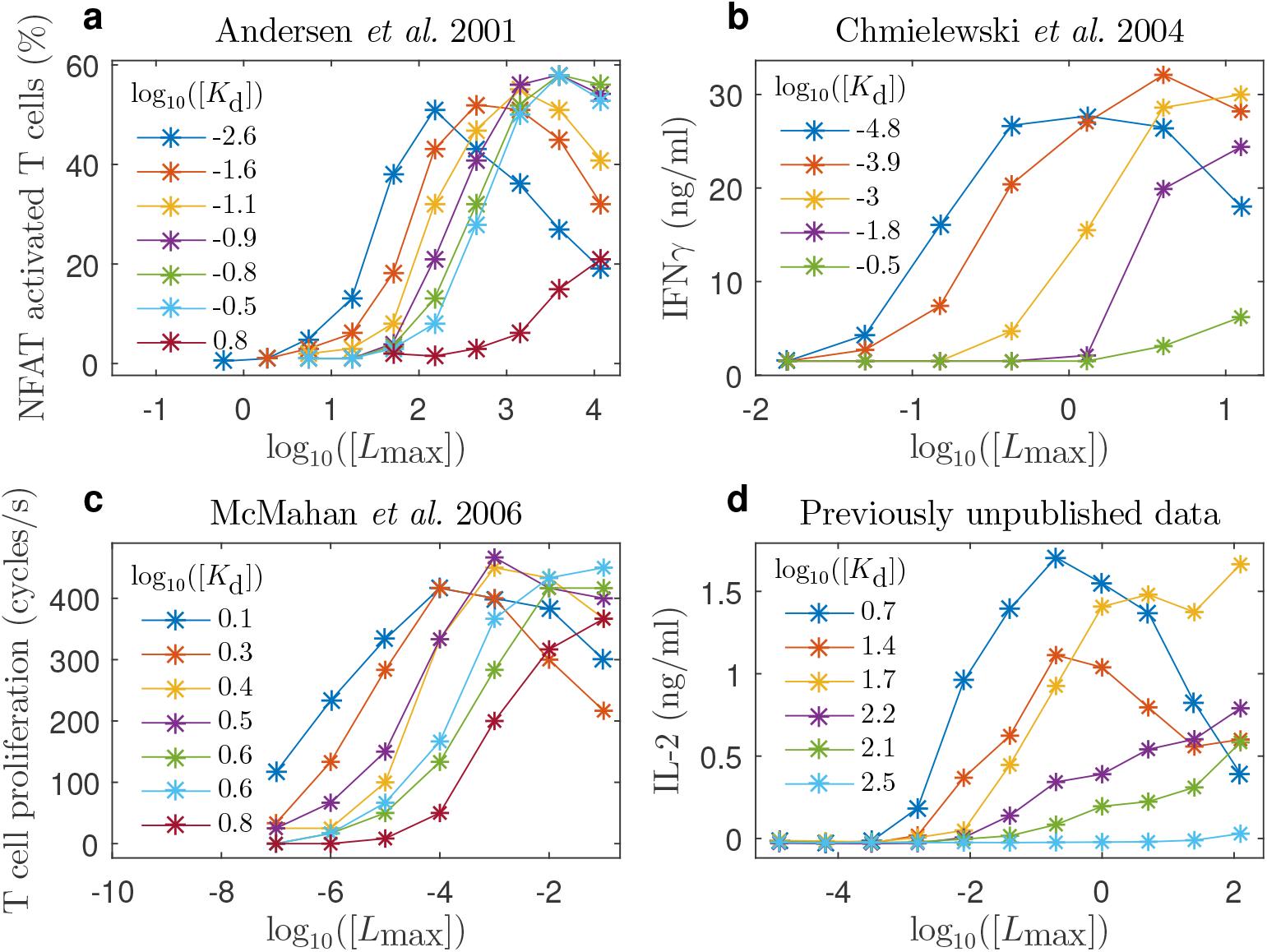
Experimental data is consistent with theory. Experimental dose-response curves (**a**) from [64] using A5 T cells expressing the 14.3.d TCR, (**b**) from [65] using T cells expressing anti-ErbB2 CARs, (**c**) from [53] using T cells expressing an AH1-specific TCR and (**d**) using naïve T cells transduced with the 1G4 TCR (see **Materials and Methods**). Dose on the x-axis is given by the concentration of (**a**) antibodies (in μm^−2^), (**b**) ErbB2 proteins (in μM) and (**c,d**) peptide (in μM). The legend gives the 3D affinity (in μM). Response on the y-axis is given by (**a**) the percentage of T cells in which the transcription factor NFAT was activated, (**b**) the density of the cytokine IFN*γ* that was produced, (**c)** the rate of T cell proliferation, and (**d**) the density of the cytokine IL-2 that was produced.

In addition, Lever *et al.* [66] summarized the results of their extensive dose-response experiments (some of which have been recently repeated [69]) in the following four statements:

1. Dose-response curves are bell-shaped for high- but not low-affinity pMHC ligands.
2. The peak amplitude of bell-shaped dose-response curves is independent of affinity.
3. A single intermediate affinity pMHC ligand produces largest response at low pMHC doses.
4. Different intermediate affinity pMHC ligands produce the largest response at high pMHC doses.

Statements 1, 2 and 4 are consistent with our model predictions shown in **Fig. 3**, as well as the other experimental dose-response data-sets shown in **Fig. 4**. However, this is not the case for statement 3, where our theory and the other dose-response data-sets show that the pMHC ligand with the highest physiological affinity produces the largest response at low doses. This discrepancy might be explained by the affinity enhanced 1G4 TCR used in the Lever *et al*. [66] (and subsequent [69]) study that extends to very high supra-physiological affinities.

Note that validating our 2D affinity based model with dose-response data-sets in which the affinity has been measured via 3D techniques (such as surface plasmon resonance) is justified because 2D affinity has been shown to correlate positively with 3D affinity, despite their respective kinetics not exhibiting such positive correlations [32, 70]. Collectively, these results indicate that simple information theoretic reasoning can help interpret complex dose-response data, and suggest that the T cell response is regulated by TCR-pMHC fluctuations.

## Discussion

A general communication system consists of at least three interconnected parts: an information source, a channel and a destination [71]. In the context of T cell activation, TCR-pMHC binding dynamics can be thought of as the information source; intracellular signaling pathways as the channel; and the cell nucleus as the destination. From this perspective, stochasticity in TCR-pMHC binding dynamics generates a ‘message’ which, depending on the kinetic rate parameters, may contain more or less information. Moreover, the average information content per second of this message, as assessed by the entropy rate, represents the average rate at which peptide-specific information is conveyed to the nucleus via signaling pathways. Based on this reasoning, we propose that T cell activation is regulated by the entropy rate of the TCR-pMHC binding dynamics. More generally this reasoning suggests that tools from information theory may help to shed light on the complex information processing mechanisms involved in T cell activation.

Indeed, a recent study by Ganti *et al.* [72] focused on channel capacity via a relatively complex model of the T cell signaling pathway. This is in contrast to our approach where we have focused on the information source via a relatively simple model of the pMHC-TCR binding dynamics. Ganti *et al.* [72] also advocate kinetic proof-reading whereby pMHC ligands are required to remain bound to TCRs for a sufficiently long time in order to initiate T cell signaling. Although the model developed here has no requirement for a minimum duration of engagement, the bio-chemical reactions of kinetic proof-reading (e.g. [5]) and reversible conformational change (Eqs. 6–9 and section 5 of the **Supplementary Information**) are similar in that both require a TCR to undergo at least one additional transition to an active (or ‘signalling-competent’ [5]) state following initial TCR-pMHC binding before a signal can be generated.

A previous stochastic model of TCR-pMHC reversible heterodimerization [73] similarly advocated that the initiation of T cell signaling required a given number of TCR-pMHC complexes to remain bound for a given minimum duration. In contrast, we posit that faster kinetics can potentially be advantageous to the T cell by increasing the entropy/signaling rate (*cf* the left and right columns of panels in **Fig. 2**). Consequently, our stochastic model is consistent with the ‘fast kinetics based serial engagement model’ [32, 1, 34], that (like our study) is informed by 2D kinetics. Furthermore, our model is in broad agreement with two recent studies that observed a temporal sequence of short-lived pMHC-TCR binding events [74, 35] that were ‘sufficiently close in space’ [74] or required ‘two or more TCRs within a range of 20 nm’ [35]. Note that in Eq. 10 an activating signal is, indeed, generated by two TCRs (one bound and one unbound) that are in close spatial proximity to each other.

Our model is also arguably in accordance with the ‘sustained signalling model’ [75, 52, 5] in that, here, TCRs remain active for a period of time after unbinding, during which they can contribute to signaling, before reverting back to their inactive (or ‘basal’ [75]) state. Moreover, this feature of our model could provide the ‘memory’ that has been suggested as necessary for conformational change models to be compatible with ‘confinement time models’ [76, 77]. In addition, our model is consistent with the ‘integrated TCR triggering model’ [15] in which TCR-pMHC binding leads to segregation of the TCR-CD3 complex from phosphatases, as well as conformational change and aggregation in the TCR-CD3 cytoplasmic tails. Thus, although we do not explicitly model phosphatase segregation, by similarly assuming that such segregation occurs upon TCR-pMHC binding then our model is arguably compatible with the ‘kinetic-segregation model’ [16, 78, 79].

Previous studies have argued that experimentally observed bell-shaped dose-responses can be explained by a TCR-proximal negative feed-back loop [80], TCR-proximal incoherent feed-forward loop [66, 67], TCR down-regulation [68], CAR down-regulation [63] or CAR dimerization [62]. Such minimal models were largely based on a deterministic framework that accounts for average copy-numbers but not their fluctuations. Although a deterministic model of serial TCR-pMHC engagement, reversible TCR conformational change and TCR aggregation gives a signaling rate similar to that shown in **Fig. 3d** (see section 5 of the **Supplementary Information**), only by taking a stochastic view have we been able to provide an information-theoretic explanation for why these three mechanisms might be utilized by the T cell.

The implications of these considerations are perhaps most important for designing the next generation of immunotherapies. For example, identifying the optimal affinity and dose is central to the design of CAR T cell therapies [81, 82, 83, 57, 56, 58, 67, 84, 63, 59] as well as cancer vaccines [53, 54, 55, 56, 60]. Our information-theoretic approach provides a framework to guide the optimization of the T cell response via modification of the affinity or dose of the TCR-pMHC binding dynamics. This issue is considered further in section 2.4 of the **Supplementary Information**, where we provide a numerical procedure to calculate the optimal affinity under conditions in which both the total number of TCRs and pMHC ligands are fixed. If it were possible to manipulate both the binding and unbinding rates then our analysis suggests that the T cell response will increase with faster kinetics, providing that the optimal affinity is maintained.

Our results also provide a note of caution. Shannon’s seminal information theorems [71] show that it is unproductive for the entropy rate of a message to exceed the communication system’s channel capacity, because the channel capacity sets an upper limit to the rate of error-free information transmission. This suggests that there is a limit to the rate at which the T cell can process information, which is set by the intracellular signaling pathways that transmit signals from the cell surface to the nucleus. Thus, there may be a limit to our ability to engineer T cell therapeutics based on manipulation of the TCR-pMHC kinetic rate parameters, unless the capacity of the signaling pathway(s) that transmit these messages can also somehow be increased. To quote Lombardi *et al.* [85] ‘*The goal in the field of communication engineering is to optimize the transference of information through channels conveniently designed*’. We speculate that the same may be true for T cell engineering.

Although we have focused on the T cell response, the simplicity of our model means that an information-theoretic perspective of receptor-ligand binding could have application to a wide range of other therapeutics. For instance, experimental evidence for binding-induced conformational change that subsequently induces aggregation and clustering not only exists for the T cell receptor [86] but has also been found for the B cell receptor [87, 88].

## Materials and Methods

### Fluctuations in the TCR-pMHC binding dynamics

In the context of the reactions of Eq. 1, let *B*_max_ and *U*_max_ denote the smaller and larger respectively of the total number of pMHC ligands, *L*_max_, and total number of TCRs, *R*_max_, given by:

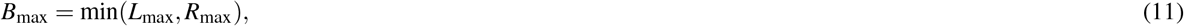

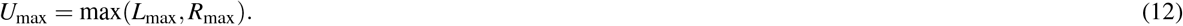

Note that *B*_max_ is also the maximum number of TCR-pMHC complexes. The stationary probability distribution of the TCR-pMHC complex number, *p*(*B*) is given by:

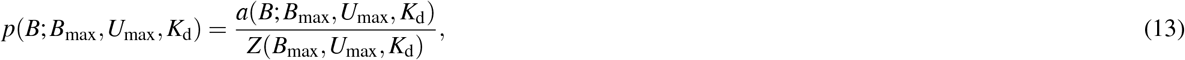

where

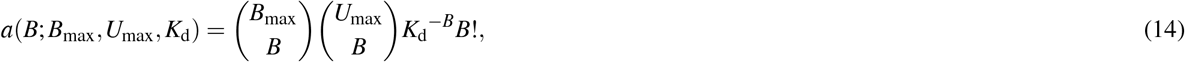

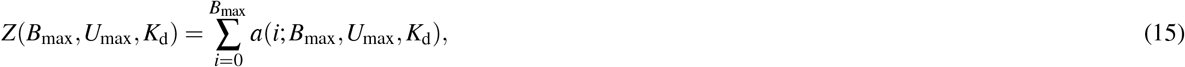

and where *K*_d_ is given by Eq. 5. The probability of there being at least one TCR-pMHC complex in the contact area, *P*_a_ is given by:

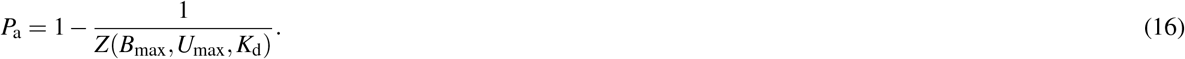

Eq. 16 is commonly referred to as the ‘probability of adhesion’ between two cells [32]. Full details of these and further calculations are provided in the **Supplementary Information**.

### Model parametrization

To produce **Figs. 2** and 3 we used parameters from the literature. Specifically, Huang *et al.* [32] performed a series of adhesion frequency assays in which a T cell was mechanically brought in and out of contact with an APC for varying durations on multiple occasions. An approximation to the probability of adhesion, *P*_a_ given by Eq. 16 (and its time-dependent generalization) was fitted to the proportion of contacts that had resulted in adhesion. Table 1 summarises the key parameters from the Huang *et al.* study which allows for conversion to the parameters described in this study (i.e. *L*_max_, *R*_max_, *K*_d_ and *k*_on_/*ν*). Combining table 1 with Eq. 5 gives an order of magnitude range of *K*_d_ ∈ [10^1^, 10^4^] and *k*_on_/*ν* ∈ [10^−4^, 10^0^]s^−1^. Table 1 also gives an order of magnitude range of *L*_max_ ∈ [10^0^, 10^2^]. We extended the upper limit of *L*_max_ by an order of magnitude because many dose-response studies consider a wider range of doses (e.g. [66]). Finally, table 1 gives a parameter estimate of *R*_max_ ~ 10^1^. Other studies have found that the number of TCRs in individual microclusters is ~ 10-100 [19, 20, 21, 32, 33]. Therefore, we also considered *R*_max_ ~ 10^2^ in **Fig. 3c** as a sensitivity analysis.

**Table 1.** Parameters derived from Huang *et al.* [32] based on experiments performed at 37 °C. **Estimation** refers to whether the parameter was directly measured or fitted from data. **Name** is how the parameter was described in the Huang *et al.* study. **Notation** denotes the parameter based on the notation in this study. **Value** gives an order of magnitude estimate or range. Note that the 2D contact area was described as ‘a few percent’ of 3 *μ*m^2^ or 1 *μ*m^2^ depending on the type of apparatus used in the experiments.

| Estimation | Name | Notation | Value | Units |
| --- | --- | --- | --- | --- |
| Measured | 2D contact area | $\nu$ | $10^{-1}$ | $\mu\text{m}^2$ |
| Measured | TCR density | $R_{\text{max}}/\nu$ | $10^2$ | $\mu\text{m}^{-2}$ |
| Measured | pMHC density | $L_{\text{max}}/\nu$ | $[10^1, 10^3]$ | $\mu\text{m}^{-2}$ |
| Fitted | Effective 2D affinity | $\nu^2/K_d$ | $[10^{-3}, 10^{-6}]$ | $\mu\text{m}^4$ |
| Fitted | 2D off rate | $k_{\text{off}}$ | $[10^0, 10^1]$ | $\text{s}^{-1}$ |

### Minimal model of signal generation

Eqs. 6, 7, 8 and 9 model a combination of the serial TCR-pMHC engagement and TCR reversible conformational change mechanisms. Specifically, Eq. 6 models an inactive TCR (i.e. a TCR in its resting state), *R*_I_ binding with a pMHC ligand, *L* to form a TCR-pMHC complex, *B*. Eq. 7 models a TCR conformational change whereby an inactive TCR enters an active state, *R*_A_, upon unbinding from the TCR-pMHC complex. For simplicity we assume that an active TCR can bind with a pMHC ligand at the same rate as an inactive TCR as shown by Eq. 8. Furthermore, Eq. 9 models an active TCR reverting to an inactive TCR. Eq. 10 models the TCR aggregation mechanism whereby a signal, *S* is generated providing that an active TCR is in sufficient proximity to a TCR-pMHC complex.

The mean signaling rate shown in **Fig. 3d** was calculated via repeated stochastic simulations of the reactions given in Eqs. 6–10 as follows. Initial conditions were: *R*_I_(0) = *R*_max_ = 10 and *R*_A_(0) = *B*(0) = *S*(0) = 0. The unbinding rate, *k*_off_ was fixed at 1/s and the binding rate, *k*_on_/*ν* was varied between 10^−4^/s and 10^−1^/s to give the values of *K*_d_ shown in the legend as calculated via Eq. 5. Each stochastic simulation was run until either *S*(*t*) > 10^4^, or *t* > 10^4^s and then the signaling rate was calculated as 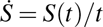. The mean signaling rate, 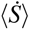 was then calculated as the mean of 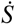 over 10 simulations for each set of parameter values.

### T cell and monocyte preparation

The assay was performed as previously detailed [89, 90]. In brief, human autologous T cells and monocytes were isolated from anonymised leukopoiesis products obtained from the NHS at Oxford University Hospitals (REC 11/H0711/7). Naïve T cells were isolated using negative selection kits (Stemcell Technologies).

T cells were cultured at 37°C, 5% CO_2_, in RPMI 1640 (Roswell Park Memorial Institute) medium supplemented with 10% FBS (Gibco), 5% penicillin-streptomycin (PenStrep, Gibco), 1x MEM non-essential amino acids solution, 20 mM HEPES, 1mM sodium pyruvate, 2 mM Glutamax and 50 μM 2-mercaptoethanol (Sigma) (all from Thermo Fisher unless stated otherwise).

T cells were resuspended at 25 × 10^6^/ml in Opti-MEM serum free medium containing mRNA for the 1G4 TCR*α*, TCR*β* and CD3*ζ* chains at 2 μg/10^6^ cells and electroporated at 300 V, 2 ms in an ECM 830 Square Wave Electroporation System (BTX).

Monocytes were enriched using a RosetteSep kit (Stemcell Technologies) and cultured at 1-2 × 10^6^/ml in 12-well plates with 1 ml of differentiation medium containing 50 ng/ml interleukin 4 (IL-4, 200–04 A, Peprotech) and 100 ng/ml granulocyte-monocyte colony stimulating factor (GM-CSF, 11343125, Immunotools) for 24 hr. For maturation the following cytokines were added for an additional 24 hr: 1 μM prostaglandin E_2_(PGE_2_, P6532, Sigma), 10 ng/ml interleukin one beta (IL-1*β*, 201-LB-025/CF, Bio-Techne), 50 ng/ml tumour necrosis factor alpha (TNF*α*, 300-01A, Peprotech) and 20 ng/ml interferon gamma (IFN*γ*, 285-IF-100/CF, Bio-Techne).

### T cell and monocyte co-culture

Monocyte-derived dendritic cells (moDCs) were pulsed with a titration of different variants of the NYE-ESO157–165 as in [90]. Loading was done for 1–2 hr at 37°C. T cells and moDCs were mixed at 1:1 ratio and incubated for 24hrs before supernatant were collected for downstream analysis.

### ELISAs

Human interleukin two (IL-2) Ready-SET Go! ELISA kit (eBioscience/Invitrogen) and Nunc MaxiSorp 96-well plates (Thermo Fisher) were used according to the manufacturer’s instructions to test appropriately diluted (commonly 4-fold) T cell supernatant for secretion of IL-2. The mean of three independent experiments is shown in **Fig. 4d**.

## Author Contributions

JRE analyzed the data and performed the theoretical research. EAS performed the experimental research. JRE and BDM contributed new analytic tools. JRE, TE and BDM designed the theoretical research and wrote the article. EAS and OD edited the article.

## Acknowledgements

Thanks to Anna Huhn and Johannes Pettmann for helpful discussions regarding the dose-response data-sets. Thanks also to Rosanna Smith and Michael Casey for helpful discussions during the early drafting stage of this manuscript.

