## Supplementary Information for "Fluctuations in TCR and pMHC interactions regulate T cell activation"

#### ABSTRACT

This **Supplementary Information** includes derivations of the key equations in the main text. In particular, section 2 provides derivations of the equations in the first section of the **Results** in the main text, as well as the equations in the **Methods**. Sections 4 and 5 provide derivations of the equations that underpin the second section of the **Results** in the main text. Although the results in sections 4 and 5 depend on those in sections 1 and 3, readers familiar with this topic may wish to skip these earlier sections.

### 1 Exact deterministic solution

Let $R$ denote a T cell receptor, $L$ denote its associated ligand and $B$ denote the bound complex that is formed when the ligand and receptor bind together. This interaction can be described as a reversible heterodimerization reaction (RHR) [1] given as follows:

$$L + R \underset{k_{\text{off}}}{\overset{k_{\text{on}}}{\rightleftharpoons}} B, \tag{1}$$

where $k_{\text{on}} \in \mathbb{R}_+$ and $k_{\text{off}} \in \mathbb{R}_+$ are the rates of binding and unbinding, respectively. By the law of mass action [2], the RHR can be described by the following reaction rate equations:

$$\frac{d[L]}{dt} = k_{\text{off}}[B] - k_{\text{on}}[L][R], \tag{2}$$

$$\frac{d[R]}{dt} = k_{\text{off}}[B] - k_{\text{on}}[L][R], \tag{3}$$

$$\frac{d[B]}{dt} = k_{\text{on}}[L][R] - k_{\text{off}}[B], \tag{4}$$

where $t$ denotes time and $[L] \in \mathbb{R}_+$, $[R] \in \mathbb{R}_+$, $[B] \in \mathbb{R}_{\geq 0}$ are the concentrations of ligand, receptor and bound complex respectively. Adding equations 2 and 4 gives:

$$\frac{d[L]}{dt} + \frac{d[B]}{dt} = 0 \tag{5}$$

$$\Rightarrow [L(t)] + [B(t)] = [L_0] + [B_0] = [L_{\text{max}}], \tag{6}$$

where $[L_0] = [L(t=0)]$, $[B_0] = [B(t=0)]$ and $[L_{\text{max}}]$ is the total concentration of ligand (both unbound and bound), which is also equal the maximum possible concentration of (unbound) ligands. Similarly,

$$[R(t)] + [B(t)] = [R_0] + [B_0] = [R_{\text{max}}], \tag{7}$$

where $[R_0] = [R(t=0)]$ and $[R_{\text{max}}]$ is the total concentration of receptor (both unbound and bound), which is also equal to the maximum possible concentration of (unbound) receptors. It is clear that the minimum possible concentration of bound complexes is zero when there are no ligands bound to any receptors. It is also evident that the maximum possible concentration of bound complexes, $[B_{\text{max}}]$ is limited by the minimum of the total concentration of ligands and receptors. This leads us to two important definitions given by:

$$[B_{\text{max}}] = \min([L_{\text{max}}], [R_{\text{max}}]), \tag{8}$$

$$[U_{\text{max}}] = \max([L_{\text{max}}], [R_{\text{max}}]). \tag{9}$$

We can now combine equations 4, 6, 7, 8 and 9 to provide a first order non-linear ordinary differential equation:

$$\frac{d[B]}{dt} = k_{\text{on}}([B_{\text{max}}] - [B])([U_{\text{max}}] - [B]) - k_{\text{off}}[B]. \tag{10}$$

The equilibrium solution, $[B_{\text{eq}}]$, to this equation is:

$$[B_{\text{eq}}] = \frac{[B_{\max}] + [U_{\max}] + [K_{\text{d}}] - \sqrt[+]{([B_{\max}] + [U_{\max}] + [K_{\text{d}}])^2 - 4[B_{\max}][U_{\max}]}}{2}, \tag{11}$$

where $[K_{\text{d}}] = k_{\text{off}}/k_{\text{on}}$ is the deterministic (or three dimensional [3]) dissociation constant. Equivalent equations for the number of ligands and receptors are given by

$$[L_{\text{eq}}] = [L_{\max}] - [B_{\text{eq}}], \tag{12}$$

$$[R_{\text{eq}}] = [R_{\max}] - [B_{\text{eq}}], \tag{13}$$

from which it is apparent that the deterministic dissociation constant can be written as [4],

$$[K_{\text{d}}] = \frac{[L_{\text{eq}}][R_{\text{eq}}]}{[B_{\text{eq}}]}. \tag{14}$$

Finally, combining equation 14 with equations 12 and 13 we find that:

$$[B_{\text{eq}}] = \frac{[L_{\max}][R_{\text{eq}}]}{[K_{\text{d}}] + [R_{\text{eq}}]} = \frac{[R_{\max}][L_{\text{eq}}]}{[K_{\text{d}}] + [L_{\text{eq}}]}. \tag{15}$$

#### 2 Exact stochastic solution

Henceforth we will use $B$ to denote both the bound complex state as well as its copy-number. The chemical master equation for the bound complex copy-number of the RHR is:

$$\dot{p}(B,t) = r(B+1)p(B+1,t) + g(B-1)p(B-1,t) - (r(B) + g(B))p(B,t) \quad \text{for } B = 1,2,\ldots,B_{\max}-1, \tag{16}$$

$$\dot{p}(0,t) = r(1)p(1,t) - g(0)p(0,t), \tag{17}$$

$$\dot{p}(B_{\max},t) = g(B_{\max}-1)p(B_{\max}-1,t) - r(B_{\max})p(B_{\max},t), \tag{18}$$

where $p(B,t)$ is the probability that there are $B$ copies at time $t$ and $B_{\max}$ is the maximum bound complex copy-number. The propensity functions for binding and unbinding, respectively, are:

$$r(B) = k_{\text{off}}B, \tag{19}$$

$$g(B) = \frac{k_{\text{on}}}{v}(B_{\max} - B)(U_{\max} - B), \tag{20}$$

where $U_{\max}$ is the maximum of the total ligand copy-number, $L_{\max}$ and total receptor copy-number, $R_{\max}$ (*cf* equation 9) and $v$ is the contact area in which the reactions can occur. Note that the unbinding rate of the RHR, $k_{\text{on}}$ (given in equation 1) has dimensions of 1/(concentration $\times$ time) and consequently $k_{\text{on}}/v$ has dimensions of 1/time [5].

##### 2.1 Stationary distribution

The stationary solution to equations 16-18 is [6, p. 141]:

$$p(B) = a(B)p(0), \tag{21}$$

where

$$p(B) = \lim_{t\to\infty} p(B,t), \tag{22}$$
$$a(B) = \frac{g(B-1)g(B-2)\cdots g(1)g(0)}{r(B)r(B-1)\cdots r(2)r(1)}. \tag{23}$$

Equation 21 determines all $p(B)$ in terms of $p(0)$ which itself is fixed by the normalisation condition,

$$p(0) = \frac{1}{1+\sum_{i=1}^{B_{\max}} a(i)}, \tag{24}$$

and thus,

$$p(B) = \frac{a(B)}{1+\sum_{i=1}^{B_{\max}} a(i)}. \tag{25}$$

In this case,

$$g(B-1)g(B-2)\cdots g(1)g(0) = \left(\frac{k_{\text{on}}}{\nu}\right)^B \frac{B_{\max}!}{(B_{\max}-B)!}\frac{U_{\max}!}{(U_{\max}-B)!}, \tag{26}$$
$$r(B)r(B-1)\cdots r(2)r(1) = k_{\text{off}}{}^B B!, \tag{27}$$

and so,

$$a(B;B_{\max},U_{\max},K_{\text{d}}) = \binom{B_{\max}}{B}\binom{U_{\max}}{B} K_{\text{d}}{}^{-B} B!, \tag{28}$$

where $K_{\text{d}} = \nu k_{\text{off}}/k_{\text{on}}$ is the stochastic (or two dimensional [3]) dissociation constant.

##### 2.2 Probability of adhesion, mean, variance and Shannon entropy

The probability of adhesion between two cells is the probability that there is at least one bound complex at any given time [7, 8],

$$P_{\text{a}} = p(B>0;B_{\max},U_{\max},K_{\text{d}}) = 1-p(B=0;B_{\max},U_{\max},K_{\text{d}}) = 1-\frac{1}{\sum_{i=0}^{B_{\max}}\binom{U_{\max}}{i}\binom{B_{\max}}{i}K_{\text{d}}{}^{-i}i!}. \tag{29}$$

The mean, $\langle B\rangle$, variance, $\text{Var}(B)$ and Shannon entropy, $H(B)$ of the stationary distribution of the bound complex number can be calculated using their definitions [6, 9] in combination with equation 25 as follows:

$$\langle B\rangle = \sum_{i=0}^{B_{\max}} ip(i) = \frac{\sum_{i=0}^{B_{\max}} ia(i)}{\sum_{i=0}^{B_{\max}} a(i)}, \tag{30}$$
$$\text{Var}(B) = \sum_{i=0}^{B_{\max}} i^2 p(i) - \left(\sum_{i=0}^{B_{\max}} ip(i)\right)^2 = \langle B^2\rangle - \langle B\rangle^2, \tag{31}$$
$$H(B) = -\sum_{i=0}^{B_{\max}} p(i)\log_2(p(i)) = \langle -\log_2(p(B))\rangle. \tag{32}$$

It is important to note that equation 32 is applicable when $p(B)$ represents a probability mass function describing a collection of random variables that are *independent* and identically distributed. However, because the RHR is a one-step stochastic process, the number of bound complexes in the immediate future *depends* on the number of bound complexes at present. While estimators for the Shannon entropy of one-step stochastic processes are currently lacking [10], it is known that equation 32 provides an upper bound of the Shannon entropy when events are not independent [11]. Henceforth, to allow analytic progress we will assume that this bound is tight. Furthermore, the Shannon entropy has units of bits but in the context of the RHR, the Shannon entropy actually represents the number of bits per reaction [12].

The variance can alternatively be given as a function of the mean as follows. The mean, $\langle B \rangle$ evolves as [6, p. 140]:

$$\frac{d\langle B \rangle}{dt} = \langle g(B) \rangle - \langle r(B) \rangle. \tag{33}$$

Substituting equations 19 and 20 into equation 33 we obtain:

$$\frac{d\langle B \rangle}{dt} = \frac{k_{\text{on}}}{v} \langle (B_{\max} - B)(U_{\max} - B) \rangle - k_{\text{off}} \langle B \rangle. \tag{34}$$

At equilibrium this yields,

$$\langle B^2 \rangle = (B_{\max} + U_{\max} + K_{\text{d}}) \langle B \rangle - B_{\max} U_{\max}, \tag{35}$$

Substituting equation 35 into equation 31 gives:

$$\text{Var}(B) = \langle B \rangle (B_{\max} + U_{\max} + K_{\text{d}} - \langle B \rangle) - B_{\max} U_{\max}. \tag{36}$$

There also exists a relationship between the Shannon entropy and the variance. It has been shown that the Shannon entropy of an integer-valued random variable $B$ with finite variance $\text{Var}(B)$ satisfies the inequality [13]:

$$H(B) < \frac{1}{2} \log_2 \left( 2\pi e \left( \text{Var}(B) + \frac{1}{12} \right) \right) \tag{37}$$

##### 2.3 Variance rate and entropy rate

Adding equations 19 and 20 gives

$$\alpha(B) = r(B) + g(B) = \frac{k_{\text{on}}}{v} B_{\max} U_{\max} + \left( k_{\text{off}} - \frac{k_{\text{on}}}{v} (B_{\max} + U_{\max}) \right) B + \frac{k_{\text{on}}}{v} B^2. \tag{38}$$

$\alpha(B)$ can interpreted in three different ways [14]. First, $\alpha(B)$ represents the probability per unit time that either a binding reaction or an unbinding reaction occurs. Second, the reciprocal of $\alpha(B)$ represents the mean of an exponential distribution which can be sampled to generate the time until the next reaction. Third, $\alpha(B)$ can be interpreted as the number of reactions occurring per unit time (i.e. the rate at which the RHR proceeds). Utilizing this third interpretation, the entropy rate, $H'(B)$ (with units of bits per second) can be calculated by multiplying the Shannon entropy by the mean reaction rate, $\langle \alpha(B) \rangle$ [12]. First, we can utilize equations 30, 31, 36 and 38 to calculate the mean reaction rate:

$$\langle \alpha(B) \rangle = \sum_{i=0}^{B_{\max}} \alpha(i) p(i) \tag{39}$$

$$= \frac{k_{\text{on}}}{v} B_{\max} U_{\max} \left( \sum_{i=0}^{B_{\max}} p(i) \right) + \left( k_{\text{off}} - \frac{k_{\text{on}}}{v} (B_{\max} + U_{\max}) \right) \left( \sum_{i=0}^{B_{\max}} i p(i) \right) + \frac{k_{\text{on}}}{v} \left( \sum_{i=0}^{B_{\max}} i^2 p(i) \right) \tag{40}$$

$$= \frac{k_{\text{on}}}{v} B_{\max} U_{\max} + \left( k_{\text{off}} - \frac{k_{\text{on}}}{v} (B_{\max} + U_{\max}) \right) \langle B \rangle + \frac{k_{\text{on}}}{v} \left( \text{Var}(B) + \langle B \rangle^2 \right) \tag{41}$$

$$= \frac{k_{\text{on}}}{v} B_{\max} U_{\max} + \left( k_{\text{off}} - \frac{k_{\text{on}}}{v} (B_{\max} + U_{\max}) \right) \langle B \rangle + \frac{k_{\text{on}}}{v} (F_{\text{B}} \langle B \rangle - B_{\max} U_{\max}) \tag{42}$$

$$= 2 k_{\text{off}} \langle B \rangle. \tag{43}$$

Second, using equation 43, the entropy rate of the stationary distribution of the bound complex number is given by:

$$H'(B) = \langle \alpha(B) \rangle H(B) = 2 k_{\text{off}} \langle B \rangle H(B). \tag{44}$$

Combining equations 37 and 44 we find that:

$$H'(B) < k_{\text{off}} \langle B \rangle \log_2 \left( 2 \pi e \left( \text{Var}(B) + \frac{1}{12} \right) \right) \tag{45}$$

Although equation 45 shows that a relationship exists between the entropy rate and the variance/mean, in order to make analytical progress in section 5, we also define the variance rate of the stationary distribution of the bound complex number, $\text{Var}'(B)$ as follows:

$$\text{Var}'(B) = \langle \alpha(B) \rangle \, \text{Var}(B) = 2 k_{\text{off}} \langle B \rangle \, \text{Var}(B). \tag{46}$$

##### 2.4 Maximum variance rate and entropy rate

In order to calculate the value of $K_{\text{d}}$ at which the variance rate and entropy rate are maximised, we progress in a number of stages. First we differentiate equation 28 with respect to $K_{\text{d}}$:

$$\frac{\partial a(B)}{\partial K_{\text{d}}} = -B (K_{\text{d}})^{-B-1} \binom{B_{\max}}{B} \binom{U_{\max}}{B} B! = \frac{-B a(B)}{K_{\text{d}}}. \tag{47}$$

We can now use equation 47 to differentiate the denominator of equation 25:

$$\frac{\partial}{\partial K_{\text{d}}} \left( \sum_{i=0}^{B_{\max}} a(i) \right) = \frac{-\sum_{i=0}^{B_{\max}} i a(i)}{K_{\text{d}}}. \tag{48}$$

Substituting equation 30 into equation 48 gives:

$$\frac{\partial}{\partial K_{\text{d}}} \left( \sum_{i=0}^{B_{\max}} a(i) \right) = \frac{-\langle B \rangle}{K_{\text{d}}} \left( \sum_{i=0}^{B_{\max}} a(i) \right). \tag{49}$$

We are now in a position to use the quotient rule to evaluate the partial derivative of $p(B)$ with respect to $K_{\mathrm{d}}$:

$$\frac{\partial p(B)}{\partial K_{\mathrm{d}}} = \frac{\frac{-Ba(B)}{K_{\mathrm{d}}}\left(\sum_{i=0}^{B_{\max}} a(i)\right) + \frac{\langle B\rangle a(B)}{K_{\mathrm{d}}}\left(\sum_{i=0}^{B_{\max}} a(i)\right)}{\left(\sum_{i=0}^{B_{\max}} a(i)\right)^2} = \frac{p(B)}{K_{\mathrm{d}}}\left(\langle B\rangle - B\right). \tag{50}$$

Combining equations 50, 30 and 36 allows us to evaluate the derivative of $\langle B\rangle$ with respect to $K_{\mathrm{d}}$:

$$\frac{\partial \langle B\rangle}{\partial K_{\mathrm{d}}} = \sum_{i=0}^{B_{\max}} i\frac{\partial p(i)}{\partial K_{\mathrm{d}}} = \frac{\langle B\rangle}{K_{\mathrm{d}}}\left(\sum_{i=0}^{B_{\max}} ip(i)\right) - \frac{1}{K_{\mathrm{d}}}\left(\sum_{i=0}^{B_{\max}} i^2 p(i)\right) = \frac{-\operatorname{Var}(B)}{K_{\mathrm{d}}}. \tag{51}$$

Next, differentiating equation 36 with respect to $K_{\mathrm{d}}$ gives:

$$\frac{\partial \operatorname{Var}(B)}{\partial K_{\mathrm{d}}} = \frac{\partial \langle B\rangle}{\partial K_{\mathrm{d}}}\left(F_{\mathrm{B}} - 2\langle B\rangle\right) + \langle B\rangle = \frac{K_{\mathrm{d}}\langle B\rangle - \operatorname{Var}(B)\left(F_{\mathrm{B}} - 2\langle B\rangle\right)}{K_{\mathrm{d}}}. \tag{52}$$

Similarly, differentiating equation 32 with respect to $K_{\mathrm{d}}$ using the product rule gives:

$$\frac{\partial H(B)}{\partial K_{\mathrm{d}}} = -\left(\sum_{i=0}^{B_{\max}} \frac{\partial p(i)}{\partial K_{\mathrm{d}}}\log_2(p(i))\right) - \frac{1}{\log(2)}\left(\sum_{i=0}^{B_{\max}} \frac{\partial p(i)}{\partial K_{\mathrm{d}}}\right), \tag{53}$$

$$= -\frac{\langle B\rangle}{K_{\mathrm{d}}}\left(\sum_{i=0}^{B_{\max}} p(i)\log_2(p(i))\right) + \frac{1}{K_{\mathrm{d}}}\left(\sum_{i=0}^{B_{\max}} ip(i)\log_2(p(i))\right), \tag{54}$$

$$= \frac{\langle B\rangle H(B) - J(B)}{K_{\mathrm{d}}}, \tag{55}$$

where,

$$J(B) = -\sum_{i=0}^{B_{\max}} ip(i)\log_2(p(i)). \tag{56}$$

Using equations 51 and 52 we can differentiate equation 46 to give:

$$\frac{\partial \operatorname{Var}'(B)}{\partial K_{\mathrm{d}}} = 2k_{\mathrm{off}}\left(\operatorname{Var}(B)\frac{\partial \langle B\rangle}{\partial K_{\mathrm{d}}} + \langle B\rangle\frac{\partial \operatorname{Var}(B)}{\partial K_{\mathrm{d}}}\right), \tag{57}$$

$$= 2k_{\mathrm{off}}\left(\langle B\rangle^2 + \frac{\partial \langle B\rangle}{\partial K_{\mathrm{d}}}\left(\operatorname{Var}(B) + \langle B\rangle(F_{\mathrm{B}} - 2\langle B\rangle)\right)\right), \tag{58}$$

$$= 2\frac{k_{\mathrm{on}}}{v}\left(K_{\mathrm{d}}\langle B\rangle^2 - \operatorname{Var}(B)\left(\operatorname{Var}(B) + \langle B\rangle(F_{\mathrm{B}} - 2\langle B\rangle)\right)\right). \tag{59}$$

Using equation 59 we find that the variance rate is maximised when:

$$K_{\mathrm{d}}\langle B\rangle^2 - \operatorname{Var}(B)\left(\operatorname{Var}(B) + \langle B\rangle(F_{\mathrm{B}} - 2\langle B\rangle)\right) = 0. \tag{60}$$

Similarly, utilizing equations 51 and 55 we can differentiate equation 44 to give:

$$\frac{\partial H'(B)}{\partial K_{\mathrm{d}}} = 2k_{\mathrm{off}}\left(H(B)\frac{\partial \langle B\rangle}{\partial K_{\mathrm{d}}} + \langle B\rangle\frac{\partial H(B)}{\partial K_{\mathrm{d}}}\right), \tag{61}$$

$$= 2\frac{k_{\mathrm{on}}}{v}\left(\langle B\rangle\left(\langle B\rangle H(B) - J(B)\right) - \operatorname{Var}(B)H(B)\right). \tag{62}$$

Using equation 62 we find that the entropy rate is maximised when:

$$\langle B\rangle\left(\langle B\rangle H(B)-J(B)\right)-\mathrm{Var}(B)H(B)=0. \tag{63}$$

For a given value of $L_{\max}$ and $R_{\max}$, the value of $K_{\mathrm{d}}$ that maximises the variance rate and entropy rate can be calculated from equations 60 and 63 using a root finding algorithm (e.g. the `fzero` function in MATLAB ®).

#### 3 Approximate deterministic solution

Consider the scenario where $[U_{\max}]$ (but not $[B_{\max}]$) is sufficiently larger than $[B(t)]$ that the larger of $[L(t)]$ and $[R(t)]$ is approximately constant. Under these conditions the second order RHR given by equation 1 is approximated by the pseudo first order reaction [15] given by:

$$L \underset{k_{\mathrm{off}}}{\overset{k_{\mathrm{on}}\,R_{\max}}{\rightleftharpoons}} B \quad \text{for } [R_{\max}] \gg [L_{\max}], \tag{64}$$
$$R \underset{k_{\mathrm{off}}}{\overset{k_{\mathrm{on}}\,L_{\max}}{\rightleftharpoons}} B \quad \text{for } [L_{\max}] \gg [R_{\max}]. \tag{65}$$

The reaction rate equations for $[L_{\max}] \gg [R_{\max}]$ are then given by:

$$\frac{d[R_{\mathrm{app}}]}{dt}=k_{\mathrm{off}}[B_{\mathrm{app}}]-k_{\mathrm{on}}[L_{\max}][R_{\mathrm{app}}], \tag{66}$$
$$\frac{d[B_{\mathrm{app}}]}{dt}=k_{\mathrm{on}}[L_{\max}][R_{\mathrm{app}}]-k_{\mathrm{off}}[B_{\mathrm{app}}], \tag{67}$$

where $[R_{\mathrm{app}}(t)]$ and $[B_{\mathrm{app}}(t)]$ are the approximate values of $[R(t)]$ and $[B(t)]$ respectively. Adding equations 66 and 67 gives:

$$\frac{d([R_{\mathrm{app}}]+[B_{\mathrm{app}}])}{dt}=0 \tag{68}$$
$$\Rightarrow [R_{\mathrm{app}}(t)]+[B_{\mathrm{app}}(t)]=[R_{\max}]. \tag{69}$$

Substituting equation 69 into equation 67 and utilizing the symmetry of the system we arrive at a linear first order ordinary differential equation:

$$\frac{d[B_{\mathrm{app}}]}{dt}=k_{\mathrm{on}}[U_{\max}][B_{\max}]-(k_{\mathrm{on}}[U_{\max}]+k_{\mathrm{off}})[B_{\mathrm{app}}]. \tag{70}$$

The approximate equilibrium solution, $[B_{\mathrm{app_{eq}}}]$, to this equation is:

$$[B_{\mathrm{app_{eq}}}]=\frac{1}{1+K_{\mathrm{d_{app}}}}[B_{\max}], \tag{71}$$

where $K_{\mathrm{d_{app}}}$ is the approximate dissociation constant given by:

$$K_{\mathrm{d_{app}}}=\frac{[K_{\mathrm{d}}]}{[U_{\max}]}=\frac{K_{\mathrm{d}}}{U_{\max}}. \tag{72}$$

Equivalent equations for the ligands and receptors are given by:

$$[L_{\mathrm{app_{eq}}}] = \frac{K_{\mathrm{d_{app}}}}{1+K_{\mathrm{d_{app}}}}[L_{\max}] \quad \text{for } [R_{\max}] \gg [L_{\max}], \tag{73}$$

$$[R_{\mathrm{app_{eq}}}] = \frac{K_{\mathrm{d_{app}}}}{1+K_{\mathrm{d_{app}}}}[R_{\max}] \quad \text{for } [L_{\max}] \gg [R_{\max}]. \tag{74}$$

#### 4 Approximate stochastic solution

Let us assume the same approximation described in section 3 applies to absolute molecule numbers rather than concentrations. Under these conditions the non-linear propensity function given by equation 20 can be approximated by the following linear propensity function:

$$g(B)_{\mathrm{app}} = \frac{k_{\mathrm{on}}}{v}U_{\max}(B_{\max}-B). \tag{75}$$

##### 4.1 Stationary distribution

In order to derive the approximate stationary distribution, $p(B)_{\mathrm{app}}$, we start by using equation 75 to give:

$$g(B-1)_{\mathrm{app}}g(B-2)_{\mathrm{app}}\cdots g(1)_{\mathrm{app}}g(0)_{\mathrm{app}} = \left(\frac{k_{\mathrm{on}}}{v}U_{\max}\right)^{B}\frac{B_{\max}!}{(B_{\max}-B)!}. \tag{76}$$

Substituting equations 27 and 76 into equation 23 gives:

$$a(B;B_{\max},K_{\mathrm{d_{app}}})_{\mathrm{app}} = \binom{B_{\max}}{B}{K_{\mathrm{d_{app}}}}^{-B}, \tag{77}$$

and thus,

$$a(B;B_{\max},\phi_{\mathrm{d_{app}}})_{\mathrm{app}} = \binom{B_{\max}}{B}{\phi_{\mathrm{d_{app}}}}^{B}(1-\phi_{\mathrm{d_{app}}})^{-B}, \tag{78}$$

where

$$\phi_{\mathrm{d_{app}}} = \frac{1}{1+K_{\mathrm{d_{app}}}} = \frac{U_{\max}}{U_{\max}+K_{\mathrm{d}}}. \tag{79}$$

Next, substituting equation 77 into equation 24 gives:

$$p(0;B_{\max},K_{\mathrm{d_{app}}})_{\mathrm{app}} = \frac{1}{1+\sum_{i=1}^{B_{\max}}\binom{B_{\max}}{i}{K_{\mathrm{d_{app}}}}^{-i}} = \frac{1}{(1+{K_{\mathrm{d_{app}}}}^{-1})^{B_{\max}}} \tag{80}$$

$$\Rightarrow p(0;B_{\max},\phi_{\mathrm{d_{app}}})_{\mathrm{app}} = \left(1-\phi_{\mathrm{d_{app}}}\right)^{B_{\max}}. \tag{81}$$

Note that equation 80 is derived by using the single variable form of the binomial formula. Finally, substituting equations 78 and 81 into equation 21 gives a compact solution for the approximate stationary distribution of the bound complex number:

$$p(B;B_{\max},\phi_{\mathrm{d_{app}}})_{\mathrm{app}} = \binom{B_{\max}}{B}{\phi_{\mathrm{d_{app}}}}^{B}(1-\phi_{\mathrm{d_{app}}})^{B_{\max}-B}, \tag{82}$$

and thus,

$$B_{\text{app}} \sim \text{Binomial}(B_{\max}, \phi_{\text{d}_{\text{app}}}). \tag{83}$$

The smaller of the number of ligands/receptors is also binomially distributed, thus:

$$L_{\text{app}} \sim \text{Binomial}(B_{\max}, 1-\phi_{\text{d}_{\text{app}}}) \quad \text{for } R_{\max} \gg L_{\max}, \tag{84}$$

$$R_{\text{app}} \sim \text{Binomial}(B_{\max}, 1-\phi_{\text{d}_{\text{app}}}) \quad \text{for } L_{\max} \gg R_{\max}. \tag{85}$$

Therefore, if the number of ligands is much greater than the number of receptors, each receptor has a probability $\phi_{\text{d}_{\text{app}}}$ of being in a bound state and a probability $1-\phi_{\text{d}_{\text{app}}}$ of being in an unbound state.

##### 4.2 Mean, variance and variance rate

The mean of the approximate stationary solution, $\langle B\rangle_{\text{app}}$, can be calculated via the parameter values of the binomial distribution given in equation 83:

$$\langle B\rangle_{\text{app}} = B_{\max}\phi_{\text{d}_{\text{app}}} = \frac{1}{1+K_{\text{d}_{\text{app}}}} B_{\max}. \tag{86}$$

Similarly, the variance of the approximate stationary distribution, $\text{Var}(B)_{\text{app}}$, can also be calculated via its parameter values given in equation 83:

$$\text{Var}(B)_{\text{app}} = B_{\max}\phi_{\text{d}_{\text{app}}}(1-\phi_{\text{d}_{\text{app}}}) = \frac{K_{\text{d}_{\text{app}}}}{(1+K_{\text{d}_{\text{app}}})^2} B_{\max}. \tag{87}$$

Combining equations 19 and 75 gives the rate at which the approximate RHR proceeds, $\alpha(B)_{\text{app}}$, as:

$$\alpha(B)_{\text{app}} = r(B) + g(B)_{\text{app}} = \frac{k_{\text{on}}}{v} B_{\max} U_{\max} + \left(k_{\text{off}} - \frac{k_{\text{on}}}{v} U_{\max}\right) B. \tag{88}$$

Using equations 86 and 88 the approximate mean reaction rate, $\langle\alpha(B)\rangle_{\text{app}}$ is therefore given by:

$$\langle\alpha(B)\rangle_{\text{app}} = \sum_{i=0}^{B_{\max}} \alpha(i)_{\text{app}} p(i)_{\text{app}} = 2k_{\text{off}}\langle B\rangle_{\text{app}}. \tag{89}$$

Combining equations 87 and 89 gives the approximate variance rate, $\text{Var}'(B)_{\text{app}}$, as:

$$\text{Var}'(B)_{\text{app}} = 2k_{\text{off}}\langle B\rangle_{\text{app}}\,\text{Var}(B)_{\text{app}} = 2k_{\text{off}}(1-\phi_{\text{d}_{\text{app}}})\phi_{\text{d}_{\text{app}}}{}^2 B_{\max}{}^2 = 2k_{\text{off}} \frac{K_{\text{d}_{\text{app}}}}{(1+K_{\text{d}_{\text{app}}})^3} B_{\max}{}^2. \tag{90}$$

For the binomial distribution it has been shown that the variance and Shannon entropy have the same orderings [16] (i.e. changing a parameter value such that the variance increases or decreases will also result in an increase or decrease, respectively, of the Shannon entropy). This relationship between the variance and Shannon entropy provides further justification for using the variance rate as an analytically convenient proxy for the entropy rate in section 5.

#### 5 Signal generation

In this section we show that two models are able to generate a signal at a rate that approximates half the variance rate of the bound complex number. These two models satisfy the following general characteristics:

1. Receptors are initially *inactive* and can become *active* following ligand binding.
2. Reversible heterodimerization are two of the five bio-chemical reactions. Unlike receptors, ligands remain unchanged by these events.
3. Two further bio-chemical reactions are irreversible, one of which describes the transition between inactive/active receptors and the other describes the transition between unbound/bound receptors.
4. In addition, the irreversible generation of a signal requires an unbound receptor and a bound receptor, one of which must be in an active state.

Although the two models share these four characteristics, it is worth noting that the generation of a signal requires at least one ligand in model 1 but at least two ligands in model 2.

##### 5.1 Model 1

This model begins with the binding of a ligand, $L$ to an inactive receptor, $R_{\mathrm{I}}$ to form a bound complex, $B$. When the ligand unbinds from the bound complex, the inactive receptor enters an active state, $R_{\mathrm{A}}$ in which it can still subsequently bind with a ligand. Active receptors can also revert to inactive receptors at a rate $k_{\mathrm{r}}$. If an active receptor and a bound complex are in sufficiently close proximity then a signal, $S$ is generated at a rate $k_{\mathrm{s}}$. This model of signal generation is given by the following bio-chemical reactions:

$$L + R_{\mathrm{I}} \xrightarrow{k_{\mathrm{on}}} B\,, \tag{91}$$

$$B \xrightarrow{k_{\mathrm{off}}} L + R_{\mathrm{A}}\,, \tag{92}$$

$$L + R_{\mathrm{A}} \xrightarrow{k_{\mathrm{on}}} B\,, \tag{93}$$

$$R_{\mathrm{A}} \xrightarrow{k_{\mathrm{r}}} R_{\mathrm{I}}\,, \tag{94}$$

$$R_{\mathrm{A}} + B \xrightarrow{k_{\mathrm{s}}} R_{\mathrm{A}} + B + S\,. \tag{95}$$

The corresponding reaction rate equations are given by:

$$\frac{d[L]}{dt} = k_{\mathrm{off}}[B] - k_{\mathrm{on}}[L]([R_{\mathrm{I}}] + [R_{\mathrm{A}}]), \tag{96}$$

$$\frac{d[R_{\mathrm{I}}]}{dt} = k_{\mathrm{r}}[R_{\mathrm{A}}] - k_{\mathrm{on}}[L][R_{\mathrm{I}}], \tag{97}$$

$$\frac{d[R_{\mathrm{A}}]}{dt} = k_{\mathrm{off}}[B] - k_{\mathrm{on}}[L][R_{\mathrm{A}}] - k_{\mathrm{r}}[R_{\mathrm{A}}], \tag{98}$$

$$\frac{d[B]}{dt} = k_{\mathrm{on}}[L]([R_{\mathrm{I}}] + [R_{\mathrm{A}}]) - k_{\mathrm{off}}[B], \tag{99}$$

$$\frac{d[S]}{dt} = k_{\mathrm{s}}[R_{\mathrm{A}}][B]. \tag{100}$$

Adding equations 97, 98 and 99 gives:

$$\frac{d}{dt}([R_{\mathrm{I}}] + [R_{\mathrm{A}}] + [B]) = 0, \tag{101}$$

and thus,

$$[R_{\mathrm{I}}(t)]+[R_{\mathrm{A}}(t)]+[B(t)]=[R_{\mathrm{I}}(0)]+[R_{\mathrm{A}}(0)]+[B(0)]=[R_{\max}]. \tag{102}$$

Adding equations 97 and 98 gives:

$$\frac{d}{dt}([R_{\mathrm{I}}]+[R_{\mathrm{A}}])=k_{\mathrm{off}}[B]-k_{\mathrm{on}}[L]([R_{\mathrm{I}}]+[R_{\mathrm{A}}]). \tag{103}$$

If we let $[R]=[R_{\mathrm{I}}]+[R_{\mathrm{A}}]$ then equations 96, 99 and 103 are equivalent to equations 2, 4 and 3, respectively. This means that the concentrations of ligands and bound complexes follow the same dynamics as the RHR, as does the concentration of the sum of the inactive and active receptors. Equation 98 can also be used to find the equilibrium solution for the active receptors:

$$[R_{\mathrm{A_{eq}}}]=\frac{[K_{\mathrm{d}}][B_{\mathrm{eq}}]}{[L_{\mathrm{eq}}]+[K_{\delta}]}, \tag{104}$$

where $[B_{\mathrm{eq}}]$ and $[L_{\mathrm{eq}}]$ are given by equations 11 and 12, respectively, and where:

$$[K_{\delta}]=k_{\mathrm{r}}/k_{\mathrm{on}}. \tag{105}$$

Combining equations 100 and 104 gives:

$$\frac{d[S]}{dt}\to\left(\frac{d[S]}{dt}\right)_{\mathrm{eq}}=[\dot{S}]_{\mathrm{eq}}=k_{\mathrm{s}}[R_{\mathrm{A_{eq}}}][B_{\mathrm{eq}}]=k_{\mathrm{s}}\frac{[K_{\mathrm{d}}][B_{\mathrm{eq}}]^2}{[L_{\mathrm{eq}}]+[K_{\delta}]}\quad \text{as } t\to\infty. \tag{106}$$

We refer to $[\dot{S}]_{\mathrm{eq}}$ in equation 106 as the deterministic signaling rate. To simplify the calculations for illustrative purposes, let us assume that $k_{\mathrm{r}}=k_{\mathrm{s}}/\nu=k_{\mathrm{off}}$. Then by utilizing equation 15, equations 104 and 106 reduce to:

$$[R_{\mathrm{A_{eq}}}]=\frac{([R_{\max}]-[B_{\mathrm{eq}}])[B_{\mathrm{eq}}]}{[R_{\max}]}, \tag{107}$$

$$[\dot{S}]_{\mathrm{eq}}=\nu k_{\mathrm{off}}\frac{([R_{\max}]-[B_{\mathrm{eq}}])[B_{\mathrm{eq}}]^2}{[R_{\max}]}. \tag{108}$$

Note that equations 107 and 108 have a similar form to the variance and variance rate given by equations 36 and 46, respectively. We can elaborate on this connection to the variance and variance rate for the special case of $[L_{\max}]\gg[R_{\max}]$ as follows. Under such conditions, and by utilizing the approximate RHR given by equation 65, the bio-chemical reactions given by equations 91, 92, 93, 94, and 95 reduce to:

$$R_{\mathrm{A}} \underset{k_{\mathrm{off}}}{\overset{k_{\mathrm{on}}\,L_{\max}}{\rightleftharpoons}} B\,, \tag{109}$$

$$R_{\mathrm{A}} \xrightarrow{k_{\mathrm{r}}} R_{\mathrm{I}} \xrightarrow{k_{\mathrm{on}}\,L_{\max}} B\,, \tag{110}$$

$$R_{\mathrm{A}}+B \xrightarrow{k_{\mathrm{s}}} R_{\mathrm{A}}+B+S\,. \tag{111}$$

The reaction rate equations for $[L_{\max}]\gg[R_{\max}]$ are then given by:

$$\frac{d[R_{\text{Iapp}}]}{dt} = k_{\text{r}}[R_{\text{Aapp}}] - k_{\text{on}}[L_{\text{max}}][R_{\text{Iapp}}], \tag{112}$$
$$\frac{d[R_{\text{Aapp}}]}{dt} = k_{\text{off}}[B_{\text{app}}] - k_{\text{on}}[L_{\text{max}}][R_{\text{Aapp}}] - k_{\text{r}}[R_{\text{Aapp}}], \tag{113}$$
$$\frac{d[B_{\text{app}}]}{dt} = k_{\text{on}}[L_{\text{max}}]([R_{\text{Iapp}}] + [R_{\text{Aapp}}]) - k_{\text{off}}[B_{\text{app}}], \tag{114}$$
$$\frac{d[S_{\text{app}}]}{dt} = k_{\text{s}}[R_{\text{Aapp}}][B_{\text{app}}]. \tag{115}$$

As for the exact solution above, the concentrations of ligands and bound complexes follow the same dynamics as the approximate RHR, as does the concentration of the sum of the inactive and active receptors. Combining equations 71 and 113 we obtain:

$$[R_{\text{Aapp}_{\text{eq}}}] = \frac{K_{\text{d}_{\text{app}}}[B_{\text{app}_{\text{eq}}}]}{1+K_{\delta_{\text{app}}}} = \left(\frac{1}{1+K_{\delta_{\text{app}}}}\right)\left(\frac{K_{\text{d}_{\text{app}}}}{1+K_{\text{d}_{\text{app}}}}\right)[R_{\text{max}}], \tag{116}$$

where:

$$K_{\delta_{\text{app}}} = \frac{k_{\text{r}}}{k_{\text{on}}[L_{\text{max}}]}. \tag{117}$$

Equations 112 and 117 can also be combined to give:

$$[R_{\text{Iapp}_{\text{eq}}}] = K_{\delta_{\text{app}}}[R_{\text{Aapp}_{\text{eq}}}] = \left(\frac{K_{\delta_{\text{app}}}}{1+K_{\delta_{\text{app}}}}\right)\left(\frac{K_{\text{d}_{\text{app}}}}{1+K_{\text{d}_{\text{app}}}}\right)[R_{\text{max}}]. \tag{118}$$

We can now combine equations 71, 115 and 116 to obtain the approximate deterministic signaling rate:

$$[\dot{S}_{\text{app}}]_{\text{eq}} = k_{\text{s}}\left(\frac{1}{1+K_{\delta_{\text{app}}}}\right)\left(\frac{1}{1+K_{\text{d}_{\text{app}}}}\right)\left(\frac{K_{\text{d}_{\text{app}}}}{1+K_{\text{d}_{\text{app}}}}\right)[R_{\text{max}}]^2. \tag{119}$$

Thus, for our simplifying assumption of $k_{\text{r}} = k_{\text{s}}/\nu = k_{\text{off}}$, and by utilizing equations 87 and 90, equations 116 and 119 reduce to:

$$[R_{\text{Aapp}_{\text{eq}}}] = \frac{K_{\text{d}_{\text{app}}}}{(1+K_{\text{d}_{\text{app}}})^2}[R_{\text{max}}] = \frac{\text{Var}(B)_{\text{app}}}{\nu}, \tag{120}$$
$$[\dot{S}_{\text{app}}]_{\text{eq}} = \nu k_{\text{off}}\frac{K_{\text{d}_{\text{app}}}}{\left(1+K_{\text{d}_{\text{app}}}\right)^3}[R_{\text{max}}]^2 = \frac{\text{Var}'(B)_{\text{app}}}{2\nu}. \tag{121}$$

Equation 121 shows that the approximate deterministic signaling rate is equal to half the approximate variance rate of the bound complex number divided by the contact area.

In the stochastic setting, as equations 109 and 110 are both first-order reactions and $R_{\text{A}}$ in equation 111 is unchanged by the reaction, it follows that for $k_{\text{r}} = k_{\text{off}}$ the approximate mean number of active receptors is equal to the approximate variance of the bound complex number:

$$\langle R_{\text{A}}\rangle_{\text{app}} = \nu[R_{\text{Aapp}_{\text{eq}}}] = \text{Var}(B)_{\text{app}}. \tag{122}$$

Furthermore, the approximate stochastic stationary distributions of the active and inactive receptors can be derived as follows. It has been shown that the stationary distribution of a closed system of first order reactions is given by a multinomial distribution [17, 18]. The bio-chemical reactions given by equations 109 and 110 are closed via the conservation equation 102 and are first order via the approximation $L_{\max} \gg R_{\max}$. In this case, the multinomial distribution reduces to a trinomial distribution because there are three states: bound complexes, active receptors and inactive receptors. The stationary distributions of these three states are given by the marginal distributions of the trinomial distribution which are themselves binomial distributions. As such, the stationary distribution of the bound complex number remains unchanged from the approximate RHR binomial distribution given by equation 83. For the active and inactive receptors, their mean values are equivalent to equations 118 and 116 multiplied by the contact area. In addition, the number of trials remains unchanged from the approximate RHR. This gives the stationary distributions for the active and inactive receptors as:

$$R_{\mathrm{Aapp}} \sim \mathrm{Binomial}(R_{\max}, (1-\phi_{\mathrm{d_{app}}})\phi_{\delta_{\mathrm{app}}}), \tag{123}$$

$$R_{\mathrm{Iapp}} \sim \mathrm{Binomial}(R_{\max}, (1-\phi_{\mathrm{d_{app}}})(1-\phi_{\delta_{\mathrm{app}}})), \tag{124}$$

where

$$\phi_{\delta_{\mathrm{app}}} = \frac{1}{1+K_{\delta_{\mathrm{app}}}}. \tag{125}$$

Note that the probability of success in equation 123 is equivalent to the probability of a receptor being unbound *and* active, and the probability of success in equation 124 is equivalent to the probability of a receptor being unbound *and* inactive. This means that $\phi_{\delta_{\mathrm{app}}}$ is the probability of a receptor being in an active state *given* that the receptor is unbound, and $1-\phi_{\delta_{\mathrm{app}}}$ is the probability of a receptor being in an inactive state *given* that the receptor is unbound (i.e. $\phi_{\delta_{\mathrm{app}}}$ and $1-\phi_{\delta_{\mathrm{app}}}$ are conditional probabilities). For $k_{\mathrm{r}} = k_{\mathrm{off}}$ equations 123 and 124 reduce to:

$$R_{\mathrm{Aapp}} \sim \mathrm{Binomial}(R_{\max}, (1-\phi_{\mathrm{d_{app}}})\phi_{\mathrm{d_{app}}}), \tag{126}$$

$$R_{\mathrm{Iapp}} \sim \mathrm{Binomial}(R_{\max}, (1-\phi_{\mathrm{d_{app}}})^2). \tag{127}$$

The stochastic signaling rate, $\dot{S}$ is dependent on the second-order reaction in equations 95 and 111, and so analytical progress is not as straightforward as with the active and inactive receptors. However, calculation of its mean (i.e. the mean signaling rate, $\langle \dot{S} \rangle$) via a repeated stochastic simulation of the biochemical reactions given by equations 91, 92, 93, 94 and 95 indicates that a similar dependency to equation 121 holds in the stochastic setting (see second section of the **Results** in the main text). Therefore, the mean signaling rate is approximately equal to half the variance rate of the bound complex number, particularly for $L_{\max} \gg R_{\max}$.

##### 5.2 Model 2

This model begins with the binding of a ligand, $L$ to a receptor, $R$ to form an *inactive* bound complex, $B_{\mathrm{I}}$. If a second ligand is in sufficiently close proximity, then the inactive bound complex can enter an *active* state, $B_{\mathrm{A}}$ at a rate $k_{\mathrm{p}}$. Both inactive and active bound complexes can unbind at the same rate to return the original ligand and receptor to their unbound states. If a receptor and an active bound complex are in sufficiently close proximity then a signal, $S$ is generated at a rate $k_{\mathrm{s}}$. This model of signal generation is given by the following bio-chemical reactions:

$$L + R \xrightarrow{k_{\text{on}}} B_{\text{I}}\,, \tag{128}$$
$$B_{\text{I}} \xrightarrow{k_{\text{off}}} L + R\,, \tag{129}$$
$$L + B_{\text{I}} \xrightarrow{k_{\text{p}}} L + B_{\text{A}}\,, \tag{130}$$
$$B_{\text{A}} \xrightarrow{k_{\text{off}}} L + R\,, \tag{131}$$
$$R + B_{\text{A}} \xrightarrow{k_{\text{s}}} R + B_{\text{A}} + S\,. \tag{132}$$

The corresponding reaction rate equations are given by:

$$\frac{d[L]}{dt} = k_{\text{off}}([B_{\text{I}}]+[B_{\text{A}}]) - k_{\text{on}}[L][R], \tag{133}$$
$$\frac{d[R]}{dt} = k_{\text{off}}([B_{\text{I}}]+[B_{\text{A}}]) - k_{\text{on}}[L][R], \tag{134}$$
$$\frac{d[B_{\text{I}}]}{dt} = k_{\text{on}}[L][R] - k_{\text{off}}[B_{\text{I}}] - k_{\text{p}}[L][B_{\text{I}}], \tag{135}$$
$$\frac{d[B_{\text{A}}]}{dt} = k_{\text{p}}[L][B_{\text{I}}] - k_{\text{off}}[B_{\text{A}}], \tag{136}$$
$$\frac{d[S]}{dt} = k_{\text{s}}[R][B_{\text{A}}]. \tag{137}$$

Adding equations 134, 135 and 136 gives:

$$\frac{d}{dt}([R]+[B_{\text{I}}]+[B_{\text{A}}]) = 0, \tag{138}$$

and thus,

$$[R(t)]+[B_{\text{I}}(t)]+[B_{\text{A}}(t)] = [R(0)]+[B_{\text{I}}(0)]+[B_{\text{A}}(0)] = [R_{\text{max}}]. \tag{139}$$

Adding equations 135 and 136 gives:

$$\frac{d}{dt}([B_{\text{I}}]+[B_{\text{A}}]) = k_{\text{on}}[L][R] - k_{\text{off}}([B_{\text{I}}]+[B_{\text{A}}]). \tag{140}$$

If we let $[B] = [B_{\text{I}}]+[B_{\text{A}}]$ then equations 133, 134 and 140 are equivalent to equations 2, 3 and 4, respectively. This means that the concentrations of ligands and receptors follow the same dynamics as the RHR, as does the concentration of the sum of the inactive and active bound complexes. Equation 135 can also be used to find the equilibrium solution for the inactive bound complexes:

$$[B_{\text{I}_{\text{eq}}}] = \frac{[K_\rho][B_{\text{eq}}]}{[L_{\text{eq}}]+[K_\rho]}, \tag{141}$$

where $[B_{\text{eq}}]$ and $[L_{\text{eq}}]$ are given by equations 11 and 12, respectively, and where:

$$[K_\rho] = k_{\text{off}}/k_{\text{p}}. \tag{142}$$

We can now combine equations 136 and 141 to find the equilibrium solution for the active bound complexes:

$$[B_{\mathrm{A_{eq}}}] = \frac{[L_{\mathrm{eq}}][B_{\mathrm{eq}}]}{[L_{\mathrm{eq}}]+[K_\rho]}. \tag{143}$$

Combining equations 14, 132 and 143 gives the deterministic signaling rate:

$$[\dot{S}]_{\mathrm{eq}} = k_{\mathrm{s}}\frac{[K_{\mathrm{d}}][B_{\mathrm{eq}}]^2}{[L_{\mathrm{eq}}]+[K_\rho]}. \tag{144}$$

To simplify the calculations for illustrative purposes, let us assume that $k_{\mathrm{p}} = k_{\mathrm{on}}$ and $k_{\mathrm{s}}/\nu = k_{\mathrm{off}}$. Then by utilizing equation 15, equation 144 reduces to:

$$[\dot{S}]_{\mathrm{eq}} = \nu k_{\mathrm{off}}\frac{([R_{\mathrm{max}}]-[B_{\mathrm{eq}}])[B_{\mathrm{eq}}]^2}{[R_{\mathrm{max}}]}. \tag{145}$$

Note that equation 145 is equivalent to equation 108 and, therefore, has a similar form to the variance rate given by equation 46. We can elaborate on this connection to the variance rate for the special case of $[L_{\mathrm{max}}] \gg [R_{\mathrm{max}}]$ as follows. Under such conditions, and by utilizing the approximate RHR given by equation 65, the bio-chemical reactions given by equations 128, 129, 130, 131 and 132 reduce to:

$$R \underset{k_{\mathrm{off}}}{\overset{k_{\mathrm{on}}\,L_{\mathrm{max}}}{\rightleftharpoons}} B_{\mathrm{I}}\,, \tag{146}$$

$$B_{\mathrm{I}} \xrightarrow{k_{\mathrm{p}}\,L_{\mathrm{max}}} B_{\mathrm{A}} \xrightarrow{k_{\mathrm{off}}} R\,, \tag{147}$$

$$R + B_{\mathrm{A}} \xrightarrow{k_{\mathrm{s}}} R + B_{\mathrm{A}} + S\,. \tag{148}$$

The reaction rate equations for $[L_{\mathrm{max}}] \gg [R_{\mathrm{max}}]$ are then given by:

$$\frac{d[R_{\mathrm{app}}]}{dt} = k_{\mathrm{off}}([B_{\mathrm{Iapp}}]+[B_{\mathrm{Aapp}}]) - k_{\mathrm{on}}[L_{\mathrm{max}}][R_{\mathrm{app}}], \tag{149}$$

$$\frac{d[B_{\mathrm{Iapp}}]}{dt} = k_{\mathrm{on}}[L_{\mathrm{max}}][R_{\mathrm{app}}] - k_{\mathrm{off}}[B_{\mathrm{Iapp}}] - k_{\mathrm{p}}[L_{\mathrm{max}}][B_{\mathrm{Iapp}}], \tag{150}$$

$$\frac{d[B_{\mathrm{Aapp}}]}{dt} = k_{\mathrm{p}}[L_{\mathrm{max}}][B_{\mathrm{Iapp}}] - k_{\mathrm{off}}[B_{\mathrm{Aapp}}], \tag{151}$$

$$\frac{d[S_{\mathrm{app}}]}{dt} = k_{\mathrm{s}}[R_{\mathrm{app}}][B_{\mathrm{Aapp}}]. \tag{152}$$

As for the exact solution above, the concentrations of ligands and receptors follow the same dynamics as the approximate RHR, as does the concentration of the sum of the inactive and active bound complexes. Combining equations 74 and 150 we obtain:

$$[B_{\mathrm{Iapp_{eq}}}] = \left(\frac{K_{\rho_{\mathrm{app}}}}{1+K_{\rho_{\mathrm{app}}}}\right)\frac{[R_{\mathrm{app_{eq}}}]}{K_{\mathrm{d_{app}}}} = \left(\frac{K_{\rho_{\mathrm{app}}}}{1+K_{\rho_{\mathrm{app}}}}\right)\left(\frac{1}{1+K_{\mathrm{d_{app}}}}\right)[R_{\mathrm{max}}], \tag{153}$$

where:

$$K_{\rho_{\mathrm{app}}} = \frac{k_{\mathrm{off}}}{k_{\mathrm{p}}[L_{\mathrm{max}}]}. \tag{154}$$

Equations 151 and 154 can also be combined to give:

$$[B_{\mathrm{Aapp_{eq}}}] = \frac{[B_{\mathrm{Iapp_{eq}}}]}{K_{\rho_{\mathrm{app}}}} = \left(\frac{1}{1+K_{\rho_{\mathrm{app}}}}\right)\left(\frac{1}{1+K_{\mathrm{d_{app}}}}\right)[R_{\max}]. \tag{155}$$

We can now combine equations 74, 152 and 155 to obtain the approximate deterministic signaling rate:

$$[\dot{S}_{\mathrm{app}}]_{\mathrm{eq}} = k_{\mathrm{s}}\left(\frac{1}{1+K_{\rho_{\mathrm{app}}}}\right)\left(\frac{1}{1+K_{\mathrm{d_{app}}}}\right)\left(\frac{K_{\mathrm{d_{app}}}}{1+K_{\mathrm{d_{app}}}}\right)[R_{\max}]^2. \tag{156}$$

Thus, for our simplifying assumptions of $k_{\mathrm{p}} = k_{\mathrm{on}}$ and $k_{\mathrm{s}}/\nu = k_{\mathrm{off}}$, and by utilizing equation 90, equation 156 reduces to:

$$[\dot{S}_{\mathrm{app}}]_{\mathrm{eq}} = \nu k_{\mathrm{off}}\frac{K_{\mathrm{d_{app}}}}{\left(1+K_{\mathrm{d_{app}}}\right)^3}[R_{\max}]^2 = \frac{\mathrm{Var}'(B)_{\mathrm{app}}}{2\nu}. \tag{157}$$

Note that equation 157 is equivalent to equation 121, both of which show that the approximate deterministic signaling rate is equal to half the approximate variance rate of the bound complex number divided by the contact area.

In the stochastic setting, we can use the same argument described in section 5.1 to find that the stationary distribution of the receptors remains unchanged from the approximate RHR binomial distribution given by equation 85. In addition, the stationary distributions of the active and inactive bound complexes are given by:

$$B_{\mathrm{Aapp}} \sim \mathrm{Binomial}(R_{\max}, \phi_{\mathrm{d_{app}}}\phi_{\rho_{\mathrm{app}}}), \tag{158}$$

$$B_{\mathrm{Iapp}} \sim \mathrm{Binomial}(R_{\max}, \phi_{\mathrm{d_{app}}}(1-\phi_{\rho_{\mathrm{app}}})), \tag{159}$$

where

$$\phi_{\rho_{\mathrm{app}}} = \frac{1}{1+K_{\rho_{\mathrm{app}}}}. \tag{160}$$

Note that the probability of success in equation 158 is equivalent to the probability of a receptor being bound *and* active, and the probability of success in equation 159 is equivalent to the probability of a receptor being bound *and* inactive. This means that $\phi_{\rho_{\mathrm{app}}}$ is the probability of a receptor being in an active state *given* that the receptor is bound, and $1-\phi_{\rho_{\mathrm{app}}}$ is the probability of a receptor being in an inactive state *given* that the receptor is bound (i.e. $\phi_{\rho_{\mathrm{app}}}$ and $1-\phi_{\rho_{\mathrm{app}}}$ are conditional probabilities). For $k_{\mathrm{p}} = k_{\mathrm{on}}$ equations 158 and 159 reduce to:

$$B_{\mathrm{Aapp}} \sim \mathrm{Binomial}(R_{\max}, {\phi_{\mathrm{d_{app}}}}^2), \tag{161}$$

$$B_{\mathrm{Iapp}} \sim \mathrm{Binomial}(R_{\max}, \phi_{\mathrm{d_{app}}}(1-\phi_{\mathrm{d_{app}}})). \tag{162}$$

Note that equation 162 is equivalent to equation 126 but equation 161 differs from equation 127. The stochastic signaling rate, $\dot{S}$ is dependent on the second-order reaction in equations 132 and 148, and so analytical progress is not as straightforward as with the active and inactive bound complexes. However, calculation of its mean (i.e. the mean signaling rate, $\langle\dot{S}\rangle$) via a repeated stochastic simulation of the biochemical reactions given by equations 128, 129, 130, 131 and 132 indicates that a similar dependency to equation 157 holds in the stochastic setting (results not shown). Therefore, as in section 5.1, the mean signaling rate is approximately equal to half the variance rate of the bound complex number, particularly for $L_{\max} \gg R_{\max}$.
